# Distinct cell death pathways induced by granzymes collectively protect against intestinal *Salmonella* infection

**DOI:** 10.1101/2021.11.07.467595

**Authors:** Amanpreet Singh Chawla, Maud Vandereyken, Maykel Arias, Llipsy Santiago, Dina Dikovskaya, Chi Nguyen, Neema Skariah, Nicolas Wenner, Natasha B. Golovchenko, Sarah J. Thomson, Edna Ondari, Marcela Garzón-Tituaña, Christopher J. Anderson, Megan Bergkessel, Jay C. D. Hinton, Karen L. Edelblum, Julian Pardo, Mahima Swamy

## Abstract

Intestinal intraepithelial T lymphocytes (IEL) constitutively express high amounts of the cytotoxic proteases Granzymes (Gzm) A and B and are therefore thought to protect the intestinal epithelium against infection by killing infected epithelial cells. However, the role of IEL granzymes in a protective immune response has yet to be demonstrated. We show that GzmA and GzmB are required to protect mice against oral, but not intravenous, infection with *Salmonella enterica* serovar Typhimurium, consistent with an intestine-specific role. IEL-intrinsic granzymes mediate the protective effects by controlling intracellular bacterial growth and aiding in cell-intrinsic pyroptotic cell death of epithelial cells. Surprisingly, we found that both granzymes play non-redundant roles. *GzmB^-/-^* mice carried significantly lower burdens of *Salmonella*, as predominant GzmA-mediated cell death effectively reduced bacterial translocation across the intestinal barrier. Conversely, in *GzmA^-/-^*mice, GzmB-driven apoptosis favored luminal *Salmonella* growth by providing nutrients, while still reducing translocation across the epithelial barrier. Together, the concerted actions of both GzmA and GzmB balance cell death mechanisms at the intestinal epithelium to provide optimal control that *Salmonella* cannot subvert.

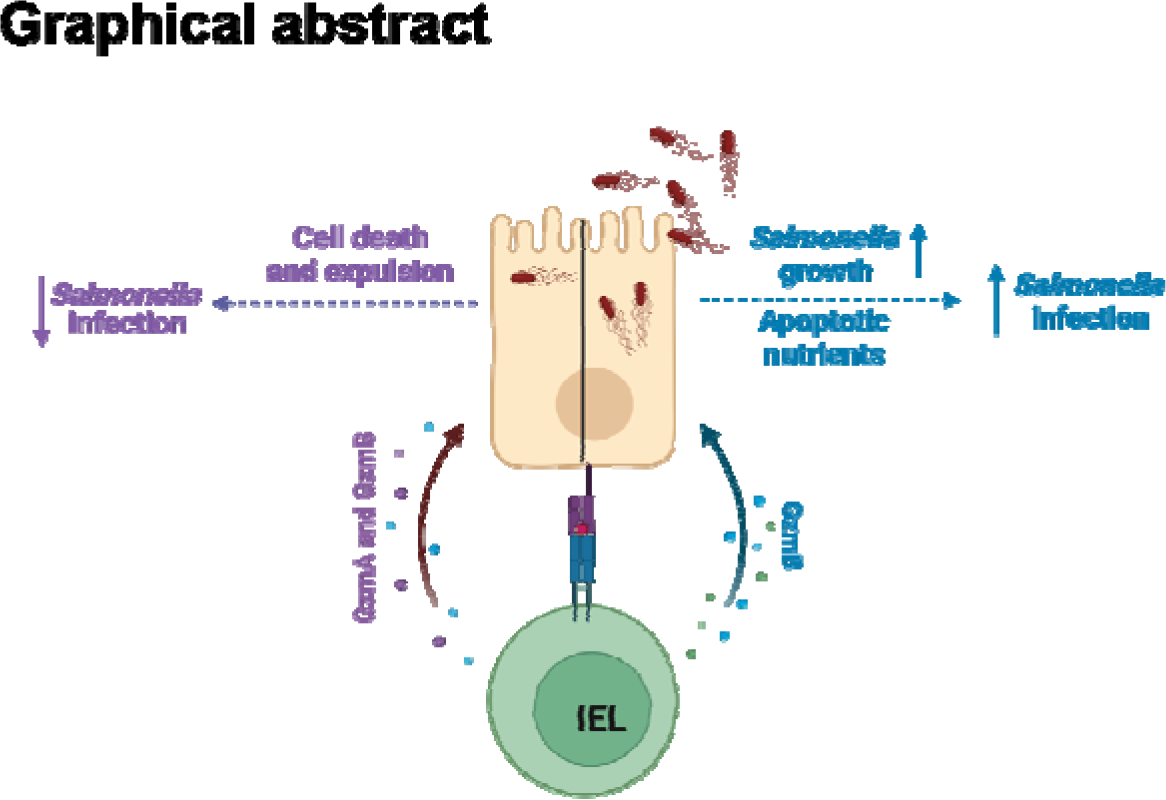

## INTRODUCTION

A key function of the intestinal epithelium is to serve as a barrier that limits the entry of microbial pathogens into the body. To protect the intestinal epithelium from such pathogens, mammals have developed sophisticated, multi-layered protective mechanisms. These include a range of innate immune defense factors produced by the intestinal epithelial cells, including mucus and antimicrobial peptides. Invasion of epithelial cells by bacterial pathogens triggers the production of pro-inflammatory cytokines that recruit immune cells to the site of infection. While the early steps of epithelial innate immune defence are well-characterized, the contributions of the different immune cells recruited to infected mucosa remain poorly understood. For example, *Salmonella enterica* serovar Typhimurium (hereafter referred to as *Salmonella* or S.Tm) infection of the mammalian intestinal epithelium induces cell death, expulsion of infected cells, inflammatory cytokine release, and activation of several types of cytotoxic lymphocytes, including natural killer cells, γδ T cells, innate lymphoid cells, mucosal-associated invariant T (MAIT) cells, yet the relative contributions of each mechanism in the control of infection are unclear^1–6^. Crucially, the role played by the cytotoxic activity of these various immune cells in controlling bacterial infection is unclear.

Cytotoxic lymphocytes kill target cells by the concerted actions of the serine proteases Granzymes (Gzm) and the pore-forming molecule Perforin, released from secretory granules at the site of contact with the target cell. Perforin mediates pore formation in the target cell membrane allowing for intracellular delivery of Gzms, where Gzms cleave critical intracellular substrates causing cell death. The human genome encodes 5 Gzms (A, B, H, K and M), whereas the mouse genome encodes 10 Gzms (A, B, C, D, E, F, G, H, K, and M)^7,8^. Granzyme A (GzmA) and B (GzmB) are the most widely studied Gzms, as they are most highly expressed in cytotoxic lymphocytes, and they are classically implicated as the effectors of granule exocytosis-mediated target cell death. GzmB induces target cell apoptosis mainly by cleaving and activating Caspase-3/7 directly or by activating mitochondrial intrinsic apoptosis pathway^9^. In contrast, GzmA can activate caspase-independent lytic cell death pathways but is less potent than GzmB. Evidence from several independent studies indicate that the key role of GzmA is to drive inflammatory responses^10^. Loss of perforin (*Prf1*) in mice abolishes granule mediated target cell death, therefore perforin is considered essential for granzyme-driven apoptosis. However, GzmA or GzmB single knockout (KO) mice do not recapitulate the loss of cytotoxic lymphocyte activity seen in Prf1 KO animals, indicating some level of redundancy between different Gzms^11^. Multiple studies have found that the individual loss of either GzmA or GzmB has no effect on the immune response to bacterial and viral infections^8^, although perforin is clearly important for protection against various viral infections^11^, and even a few bacterial pathogens^12,13^. Interestingly, extracellular levels of GzmA and GzmB were found to be increased in blood of humans infected with *Salmonella enterica* serovar Typhi^14^. Recent evidence suggests that both GzmA and GzmB have significant non-cytotoxic extracellular functions that involve cleavage of extracellular and intracellular proteins to promote wound healing, cytokine activation, inflammatory responses, and extracellular matrix remodeling 10,15–17.

One subset of T lymphocytes that constitutively express GzmA and GzmB are intestinal intraepithelial T lymphocytes (IEL)^18,19^. IEL are a heterogenous population of T cells that occupy the intercellular space between intestinal epithelial cells. IEL are classified into two main subpopulations based on ontogeny and surface receptor expression. Induced IEL consist of conventional TCRαβ CD8αβ T cells while natural IEL, characterized by the expression of CD8αα, are defined as TCRαβ CD8αα or TCRγδ CD8αα T cells. Despite developmental differences, both types of IEL are characterized by an activated phenotype that includes very high expression of GzmA and B, and lower expression of GzmC and GzmK^18^. Based on the tissue location and Gzm expression, IEL have been postulated to play a key role in defending against intestinal infection by killing infected epithelial cells. One subtype, TCRγδ IEL, have been implicated in the defense against oral *S.Tm* infection^5^ and prevent transmigration of bacteria and parasites from the gut by maintaining intestinal barrier integrity^20,21^. Changes in γδ IEL movement have been observed following infection with *Salmonella* and *Toxoplasma gondii* ^21,22^. IEL also protect against the intracellular protozoan parasite *Eimeria vermiformis*^23^. Taken together, the available data are consistent with IEL being the first immune responders to enteric pathogens. However, the exact mechanism by which IEL protect against foodborne pathogens has remained opaque, and it is currently unknown whether IEL protective functions are mediated by Gzms.

Here we address a fundamental question: Are IEL-derived Gzms pivotal for protection against foodborne intracellular bacterial pathogens such as *Salmonella* spp? Previous studies using KO mice did not find a role for Perforin and GzmB in *Salmonella* pathogenesis^24^, but these experiments involved intravenous infections rather than the oral route of *Salmonella* infection, which is the more common route of human exposure. We reveal that GzmA and GzmB act together to defend the small intestinal epithelium against oral infection. The majority of GzmA and GzmB in the gut is expressed by IELs, and our data suggest that GzmA/B are responsible for the protective effects of IEL. Surprisingly, we discovered that in the absence of GzmA, GzmB promotes bacterial growth by providing apoptosis-derived nutrients, whereas in the absence of GzmB, GzmA is more effective in preventing bacterial translocation across the intestinal barrier. Our findings reveal novel effector functions of Gzms in tissue-specific control of pathogens and explain how IEL contribute to intestinal protection.

## RESULTS

### Loss of granzymes does not affect IEL homeostasis

IEL constitutively express high levels of both GzmA and GzmB, with ∼20 million molecules of GzmA and ∼5 million molecules of GzmB per cell^18^. Flow cytometric analyses of spleen, mLNs, small intestinal (SI) and large intestinal epithelium and lamina propria (LP) layers indicated that only the SI epithelium harbors a large population of GzmA/B expressing cells **(Fig. 1A-B)**. This finding was confirmed by immunofluorescence (IF) imaging of GzmA and GzmB in the gut, that showed overlapping expression of GzmA and GzmB mainly within small round cells just below intestinal epithelial cells, but above the basement membrane (**Fig. 1C)**. Specificities of antibodies for GzmA and GzmB for both IF and flow cytometry were confirmed using *GzmA^-/-^* and *GzmB^-/-^*mice **(Suppl. Fig. 1A, E)**. These Gzm-expressing cells within the epithelium were T cells, as cells that stained positive for GzmB were also labelled with anti-CD3 **(Fig. 1D)**. Indeed, in the intestinal epithelial compartment, greater than 90% of GzmA/B+ cells are T cells, either TCRαβ^+^ or TCRγδ^+^ **(Fig. 1E)**. We conclude that the major expressers of both GzmA and GzmB in the SI are IEL.

**Figure 1.**
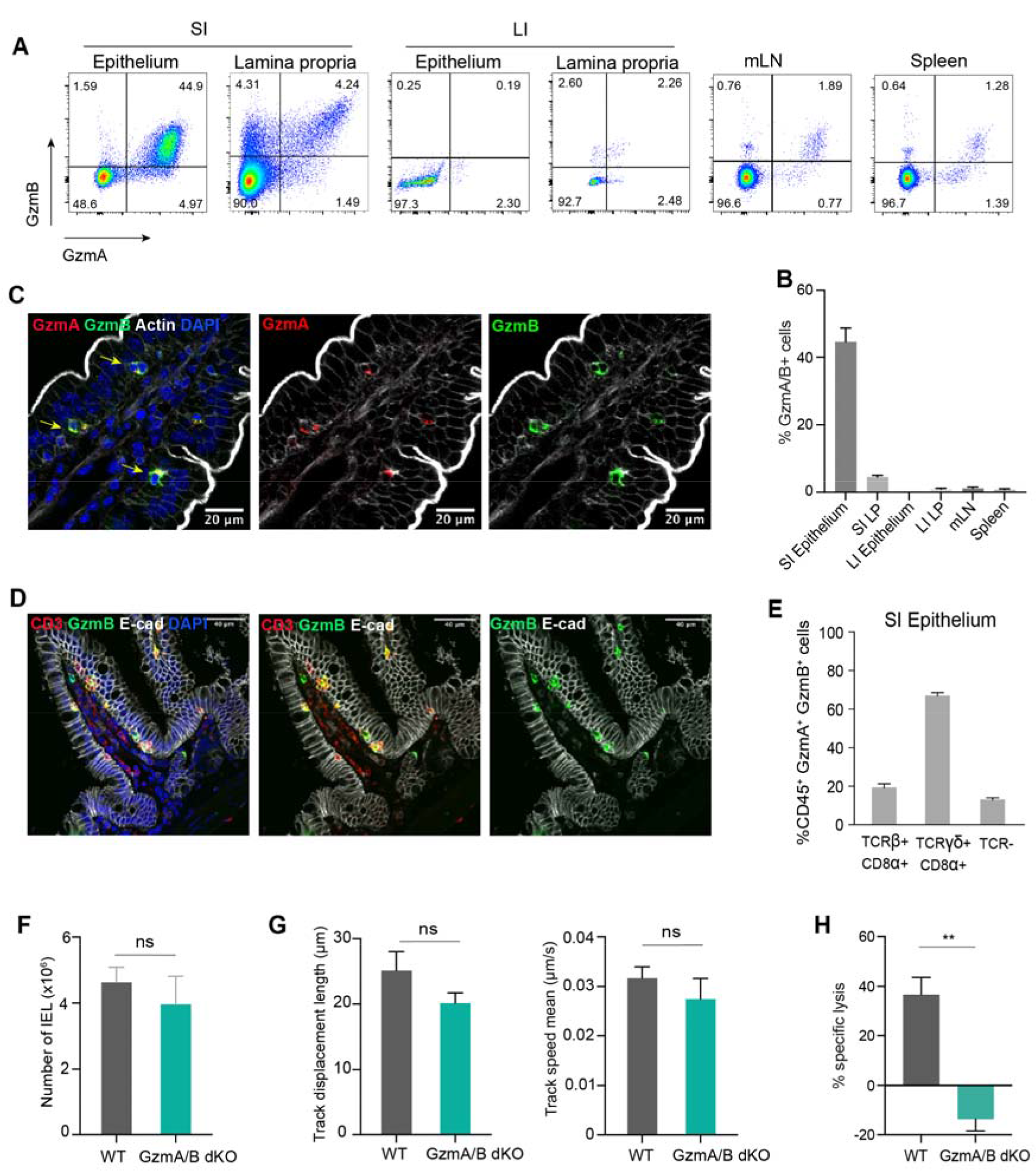
**IEL are the main source of Gzms in the mouse at steady state**. **A.** Representative flow cytometry dot plots of GzmA and GzmB expression in the small intestinal (SI) epithelium and lamina propria, large intestinal (LI) epithelium and lamina propria, mesenteric lymph nodes (mLN) and spleen of naïve WT mice (left). **B.** Bar graphs showing the quantification of GzmA^+^/GzmB^+^ cells in different organs shown in (A), n>3 mice per organ. **C.** Immunofluorescent micrographs showing cells expressing GzmA (red) and GzmB (green) in the epithelial layer of a jejunal villus. Sections were counterstained with phalloidin to show actin (white) and DAPI (blue) to show nuclei. Scale bar=20µm. **D.** Immunofluorescent micrographs showing cells expressing CD3 (red), GzmB (green) and E-cad (white) in the jejunal villus. Scale bar=40µm. **E.** Bar graphs showing which CD45^+^ cell subsets express GzmA and GzmB in the epithelium (n= 5). E-G. Bar graphs showing **F.** CD45^+^ CD103^+^ total IEL in co-housed WT and GzmA/B dKO (n= 5), **G.** Track displacement (left) and mean track speed (right) of WT and GzmA/B dKO IEL from mice which have been cohoused, in coculture with WT intestinal organoids (n=3/group). **H.** Specific cell lysis of K562 cells by either WT or GzmA/B dKO IEL from mice that have been cohoused. IEL were cultured with K562 at a 40:1 ratio, in the presence of 20ng/ml IL-15 and 1μg/ml anti-CD3 (n=3 per genotype). All data are presented as mean ± SEM. For (E-G), unpaired t-test was used to calculate the significance. Standard annotations were used to denote significance: ns: not significant, ** p<0.01.

Since IEL constitutively express Gzms, we first determined whether loss of Gzms affected IEL homeostasis. Total IEL and all subsets were present at similar numbers to wild-type (WT) mice in *GzmA^-/-^/GzmB^-/-^*(GzmA/B dKO), *GzmA^-/-^* (GzmA KO) *GzmB^-/-^* (GzmB KO) SI epithelium **(Fig. 1F, Suppl. Fig. 1B)**. Further, expression of key surface proteins that regulate IEL tissue residency (CD103, CD69), as well as recently described inhibitory receptors (CD160, LAG3, LILRB4, TIGIT, CD96) were all expressed comparably in GzmA/B dKO mice and WT mice **(Suppl. Fig. 1C)**. IEL are highly motile cells that exhibit patrolling immunosurveillance of the gut epithelium, both *in vivo* and *in vitro* ^22,25,26^. As Gzms can cleave ECM proteins, it is possible that IEL require Gzms to move efficiently within the epithelial layer. To test this, we utilized co-cultures of IEL with 3D intestinal epithelial organoids. The movement of WT and dKO IEL was tracked within the epithelial layer of co-cultures. IEL were indeed motile, with a velocity of ∼30nm.s^-1^, and both WT and GzmA/B dKO IEL moved at a similar speed **(Fig. 1G, Suppl. videos 1 and 2**). However, in the absence of GzmA/B, IEL were no longer able to kill a target cell line, K562, upon anti-CD3-mediated TCR triggering (**Fig. 1H**). Thus, except for an inability to kill target cells *in vitro*, GzmA/B dKO IEL appear to be functionally and phenotypically normal in uninfected mice.

### IEL-derived granzymes protect the intestinal epithelium against *Salmonella* infection

Since IEL are the major cell type expressing GzmA and GzmB constitutively in the mouse, we could use GzmA/B dKO to address the function of IEL-derived Gzms in intestinal infection. We infected GzmA/B dKO mice with *Salmonella enterica* Typhimurium strain SL1344 expressing GFP either orally or intravenously. We focused on mechanisms invoked during infection of the small intestinal epithelium, where IEL are present at high numbers. Hence, we chose not to pre-treat the mice with Streptomycin before infection, as Streptomycin pre-treatment increases *S*.Tm infection at the cecum and colon by ablating intestinal microbiota^27^. Indeed, when we orally gavaged C57BL6/J mice with *Salmonella* (10^9^ colony-forming units (cfu) per mouse), all mice were found to be susceptible to infection, as previously shown^28^, and *Salmonella* cfu were mainly found in the small intestine, but not in the cecum or colon (**Suppl. Fig. 2A**). Using this infection model, we found that in the absence of GzmA/B, mice were significantly more susceptible to oral *Salmonella* challenge, the natural route of infection. GzmA/B dKO displayed increased weight loss and higher bacterial loads in their mesenteric lymph nodes (mLNs), spleen, and liver (**Fig. 2A-B**). In contrast, absence of GzmA/B did not lead to increased susceptibility towards intravenous *S.*Tm infection, quantified in terms of bacterial burden and weight loss (**Fig. 2C-D**). Thus, GzmA and B contribute to protection against oral, but not intravenous infection with *Salmonella*.

**Figure 2.**
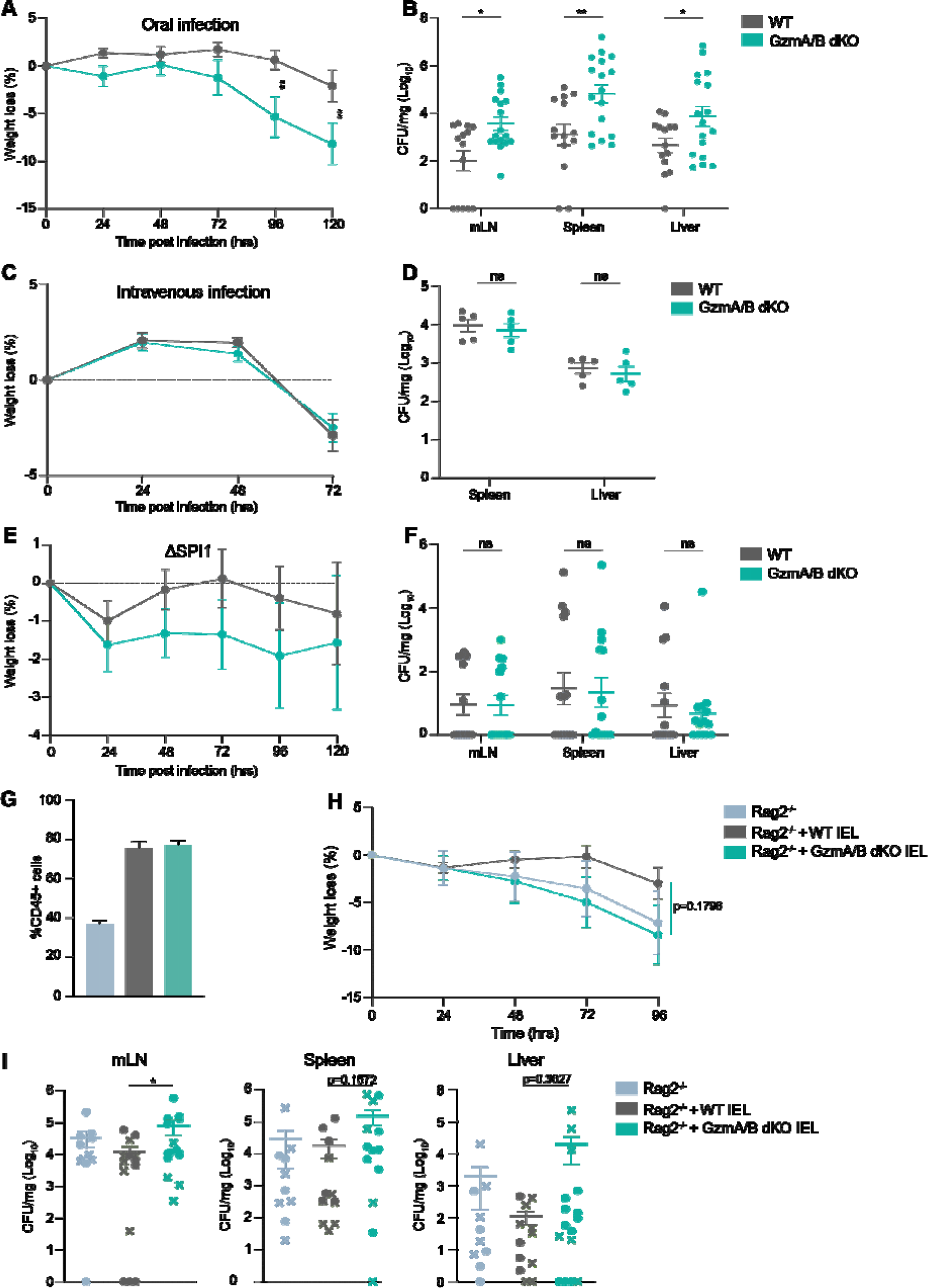
**Granzymes are important for protection against oral intestinal infection**. **A-B**. Cohoused WT (n=14) and GzmA/B dKO (n=17) mice were orally infected with SL1344-GFP and culled 5 days post infection (dpi). Weight loss (A) and CFU/mg in mLN, spleen and liver at the time of sacrifice (B) are shown. Data were pooled from 3 independent experiments. **C-D.** Cohoused WT and GzmA/B dKO mice were infected i.v. with SL1344-GFP and culled 3 dpi. Weight loss (C) and CFU/mg in spleen, liver at the time of sacrifice (D) are shown (n= 5/group). **E-F**. Cohoused WT and GzmA/B dKO mice were orally infected with ΔSPI1-SL1344 and culled 5dpi. Weight loss (E) and CFU/mg in mLN, spleen and liver are shown (F). Data were pooled from 2 independent experiments (n=14/group). **G-I**. *Rag2^-/-^* mice were adoptively transferred with either WT (n=12) or GzmA/B dKO IEL (n=15) from cohoused mice, or no IEL (n=10). 4 weeks after IEL transfer mice were orally infected with SL1344-GFP. **(G)** Percentages of CD45+ cells out of total live cells isolated from the intestinal epithelia of *Rag2^-/-^* mice, *Rag2^-/-^* mice reconstituted with WT IEL, and *Rag2^-/-^* mice reconstituted with GzmA/B dKO IEL after infection. Data were pooled from two independent experiments, where infected mice were either culled 4 or 5 dpi, depending on the severity of symptoms. Weight loss **(H)** and CFU/mg **(I)** in mLN (left), spleen (middle) and liver (right) are shown. In (I), cfu data from the two independent experiments culled 4dpi (circles) or 5dpi (crosses) are indicated. All data are presented as mean ± SEM. For bacterial counts, ranks were compared using the Mann-Whitney U-test. Standard annotations were used to denote significance: ns: not significant, * p<0.05, ** p<0.01, *** p<0.001.

*Salmonella* mainly infects the host by actively invading epithelial cells (IEC), by uptake by M cells or by phagocytic sampling of the intestinal lumen^29^. Active IEC invasion, but not the other two routes of entry, requires *Salmonella* type III secretion system 1 (TTSS-1), which is encoded by *Salmonella* pathogenicity island 1 (SPI1). To test whether epithelial invasion is necessary to activate the protective effects of Gzms, we used a ΔSPI1 strain of *Salmonella* that can still infect mice through phagocytes. We found no significant difference in the susceptibility of WT and GzmA/B dKO mice to the ΔSPI1 mutant (**Fig. 2E-F**). The levels of infection seen were low, but dKO mice also had low bacterial burdens. In summary, these data indicate that GzmA/B specifically protect small intestinal epithelial cells from *Salmonella* infection, showing that the route and site of infection are important factors in the immune response to pathogenesis.

To evaluate if IEL were the main cells mediating the protective effects of Gzms against *Salmonella* infection, we adoptively transferred WT or GzmA/B dKO IEL to *Rag2^-/-^*(RAG2 KO) mice. To improve the transfer efficiency and to enhance survival, we first cultured cells for 24h with IL-15 and retinoic acid, as we found that this combination increased the expression of the gut-homing proteins CCR9 and α4β7 **(Suppl. Fig. 2B)** and increased the efficiency of the transfer **(Suppl. Fig. 2C)**. In a competitive transfer, IEL from either WT or dKO genotypes repopulated the intestinal epithelial compartment of RAG2 KO mice equally efficiently **(Suppl. Fig. 2D)**. We next individually transferred WT or GzmA/B dKO IEL to different groups of RAG2 KO mice. After 4 weeks to allow the adoptively transferred IEL to sufficiently repopulate the gut, the mice were infected orally with *Salmonella*. Both WT and GzmA/B dKO IEL equivalently repopulated the guts of RAG2 KO mice (**Fig. 2G**). Despite large variability in the experiment, we found that overall, RAG2 KO mice reconstituted with GzmA/B dKO IEL lost more weight and had a higher infection burden than RAG2 KO with WT IEL (**Fig. 2H-I**). Moreover, three RAG2 KO that did not receive any IEL had to be culled early due to the severity to the infection, even though other Gzm-expressing innate cells had expanded to fill the epithelial niche in RAG2 KO mice (**Suppl. Fig. 2E**). These data support the hypothesis that IEL-derived Gzms are important for protection against intestinal *Salmonella* infection. Thus, the route of infection and IEL dictate the protective effects of Gzms against oral *Salmonella* infection.

### IEL utilize granzymes to kill infected epithelial cells independent of perforin

We next explored the mechanisms by which GzmA/B protect the intestinal epithelium. Gzms can cleave extracellular matrix proteins and epithelial cell junction proteins^30–32^, and loss of this activity may affect the intestinal barrier, thus increasing bacterial translocation. However, we did not find any difference in intestinal permeability to a small molecule, FITC-Dextran, in GzmA/B dKO mice compared to WT mice **(Suppl. Fig. 3A)**. Both GzmA and GzmB can regulate inflammatory cytokine production, with GzmA in particular activating key innate defense cytokines such as IL-1β, TNF or IL-6 ^15,33–36^. Therefore, the levels of cytokines in the plasma of orally infected WT and GzmA/B dKO mice were determined at day 5 post infection. We found that the levels of cytokines and chemokines in the serum of infected GzmA/B dKO mice were mostly increased compared to WT mice, especially IL-18 **(Fig. 3A)**. In intestinal ileal tissue from infected WT and GzmA/B dKO mice, there was no difference in the cytokines induced by *Salmonella* at the mRNA level **(Suppl. Fig. 3B)**. We conclude that the cytokine levels in the plasma of infected mice seem to be linked to bacterial burden rather than with the functions of Gzms in activating cytokines.

**Figure 3.**
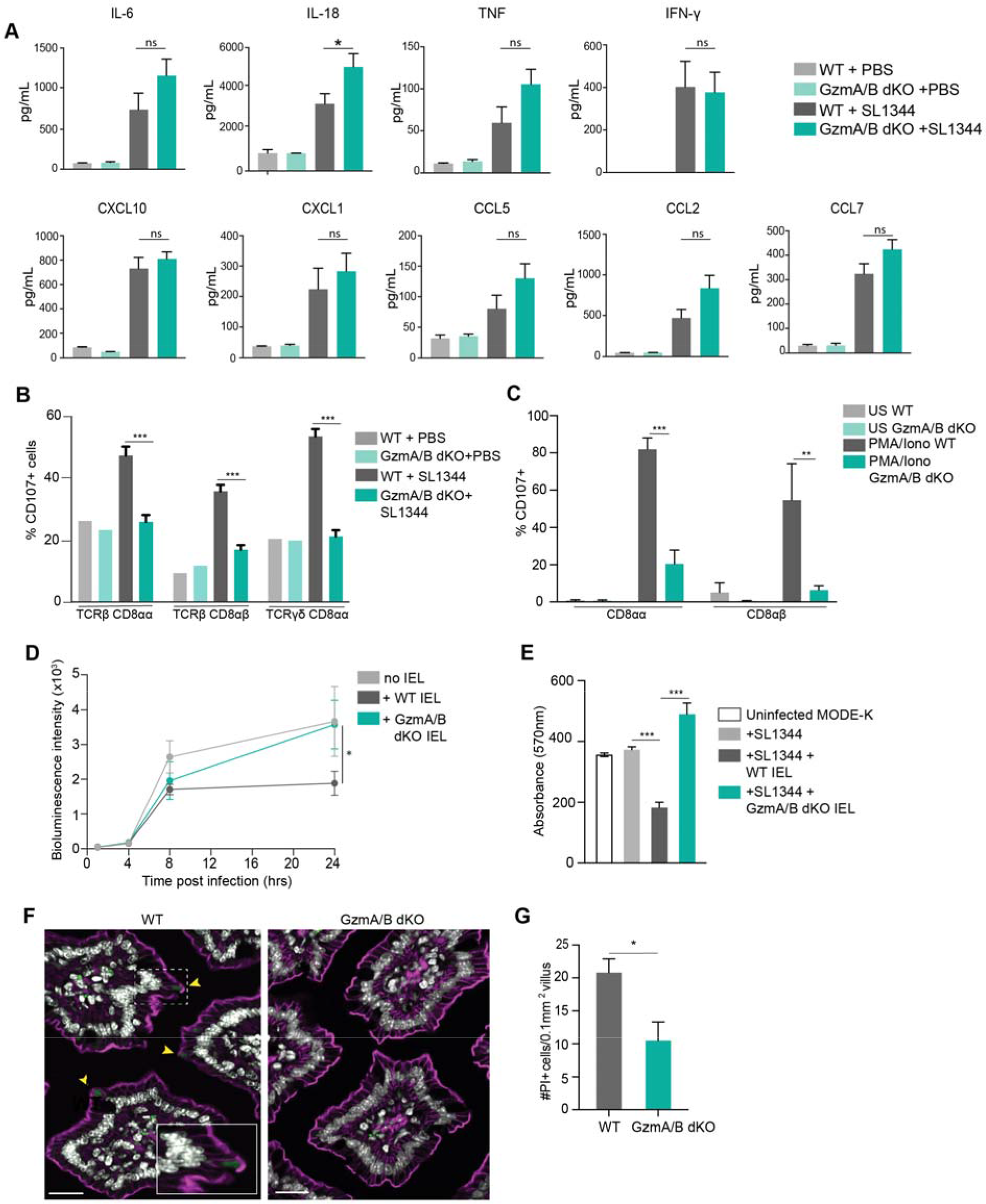
IEL use granzymes to kill infected epithelial cells. A. Chemokine and cytokine levels in plasma of naïve and orally infected cohoused GzmA/B dKO (n=9) compared to WT (n=9) from figure 2, at 5 dpi. Data were pooled from 2 independent experiments. **B.** Bar graphs showing percentage of CD107^+^ cells in co-housed WT and GzmA/B dKO IEL 5 dpi of *Salmonella* infection (n=5). **C.** Percentage of CD107^+^ cells in cultured WT and GzmA/B dKO IEL isolated from co-housed mice (n=3 each), 3h after *in vitro* PMA/ionomycin treatment. **D.** MODE-K cells were infected with SL1344-lux for 1h, treated with gentamycin, and then incubated with WT or GzmA/B dKO IEL from co-housed mice (n=3) (See Suppl. Fig. 3C for schematic). Bioluminescence intensity was measured when IEL were added, then after 4h, 8h and 24h. **E.** Infected MODE-K cells were incubated with WT or GzmA/B dKO IEL, as in (D) and infected MODE-K cells were stained with crystal violet 24h after incubation with IEL from WT and GzmA/B dKO pooled from 3 independent experiments. Absorbance of crystal violet (at 570 nm) is shown as representative of live cells. **F.** Representative images of PI^+^ enterocytes in jejunal villi of WT or GzmA/B dKO mice 1h after FlaTox treatment (0.16ug/g PA, 0.08µg/g LFn-Fla). Nuclei are shown in white PI in green and F-actin in magenta. Yellow arrowheads indicate PI^+^ enterocytes. Scale bar = 20µm, inset: scale bar = 10µm. **G.** Morphometric analysis of PI^+^ enterocytes (n=5). All data are represented as mean ± SEM. P values were calculated by (A-E) ordinary one-way ANOVA with Sidak’s multiple comparisons, (G) unpaired t-test. Standard annotations were used to denote significance: ns: not significant, * p<0.05, ** p<0.01, *** p<0.001.

The release of Gzms from intracellular secretory granules occurs through degranulation, which can be detected by the exposure of lysosome-associated membrane protein (LAMP-1 or CD107) on the surface of degranulating cells. After *Salmonella* infection, we found that over 40% of WT IEL had degranulated as determined by CD107 surface staining, indicating robust activation of IEL **(Fig. 3B)**. However, no degranulation was observed in dKO IEL from *Salmonella*-infected dKO mice. The inability of GzmAB dKO IEL to degranulate appears to be a cell-intrinsic defect, as *in vitro* after stimulation with phorbol ester PMA and calcium ionophore ionomycin, we find that WT IEL degranulate but GzmA/B dKO IEL do not **(Fig. 3C).** Because the cytotoxic functions of Gzms are generally perforin-dependent and perforin is a component of cytotoxic granules^37,38^, we asked whether perforin was required for GzmA/B mediated protection against intestinal infection. Surprisingly, we found that *Prf1^-/-^* (Pfn KO) mice were not more susceptible to oral *Salmonella* infection than WT mice (**Suppl. Fig. 4**). These data indicate that IEL utilize Gzms in a degranulation-dependent, but perforin-independent manner to control *Salmonella* infection.

To further explore how IEL and Gzms are protecting intestinal epithelial cells against infection, we utilized a cell culture infection system (**Suppl. Fig. 3C**). MODE-K cells, small intestinal epithelial cell line^39^, were first infected with a luciferase-expressing strain of *Salmonella* (SL1344-lux) for 1h, then any remaining extracellular bacteria were killed using gentamycin. At this point, purified IEL were added, and the growth of intracellular *Salmonella* was measured at various timepoints using luciferase activity as a readout (**Suppl. Fig. 3D**). We found that after 24h incubation of infected IEC with WT IEL, there was a marked reduction in *Salmonella* levels. Strikingly, no reduction in intracellular levels of SL1344-lux was seen upon co-culture with GzmA/B dKO IEL **(Fig. 3D**). Next, we investigated whether WT and GzmA/B dKO IEL were killing the infected MODE-K cells, by staining living cells with Crystal Violet **(Fig. 3E**). While incubation with WT IEL reduced the viability of MODE-K cells, GzmA/B dKO IEL did not affect the viability of the infected MODE-K cells. Note that IEL do not kill uninfected MODE-K cells (**Suppl. Fig. 3E**). Together, these data indicate that IEL specifically kill infected epithelial cells using Gzms.

### Granzymes aid in intestinal epithelial lytic cell death

It is increasingly recognized that *Salmonella* infection of the intestinal epithelium leads to activation of multiple cell death pathways^6^. *S.*Tm express flagella, type III secretion systems, LPS, and other pathogen-associated molecular patterns that rapidly activate innate immune mechanisms, including the NLRC4 inflammasome in the infected intestinal epithelial cell^6^. Activation of the NLRC4 inflammasome eventually leads to pyroptosis and expulsion of infected epithelial cells^40^, while LPS triggers apoptosis, most likely through a TNF-dependent mechanism. Previous work has shown that fewer apoptotic cells (cleaved caspase-3-positive) were seen in the intestinal epithelium of GzmA/B dKO mice treated with LPS or TNF, compared to WT mice^41^, suggesting that IEL-derived GzmA/B contribute to the induction of apoptosis under these conditions. However, the contribution of GzmA/B to *Salmonella-* induced epithelial pyroptosis is unknown. Since *Salmonella* infection events in the small intestine occur at very low frequencies and are hard to detect, we used FlaTox, an engineered form of flagellin that allows the cytosolic delivery of flagellin. FlaTox mimics *Salmonella* infection as it activates the NAIP/NLRC4 inflammasome, the cytosolic sensors for bacterial flagellin and type III secretion systems, and induces pyroptosis and expulsion of intestinal epithelial cells^40,42^. To test whether inflammasome induced pyroptosis was also reduced in the absence of Gzms, WT and GzmA/B dKO mice were treated with FlaTox for 1h, and propidium iodide (PI) to label dying cells, prior to euthanasia. Significantly fewer dying epithelial cells (PI^+^) were detected in ilea of GzmA/B dKO mice relative to WT **(Fig. 3F-G).** These data, in conjunction with the previous report on the shedding of apoptotic enterocytes^41^ indicate that Gzms contribute to epithelial cell death, even when external stimuli such as flagellin or LPS are activating cell-intrinsic death. Taken together, we conclude that IEL kill *Salmonella* infected intestinal epithelial cells via release of Gzms A and B, which together act to reduce dissemination of infection from the gut.

### GzmA and GzmB drive divergent mechanisms against intestinal infection

Our findings thus far prompted us to test the relative importance of the individual Gzms, as GzmA and GzmB are proteases with very different specificities and induce different types of cell death^11^. Upon oral *Salmonella* infection, GzmA KO mice suffered more weight loss and marginally higher bacterial burdens than WT mice (**Fig. 4A-B**). No difference in the induction of serum cytokines was seen in infected GzmA KO as compared to the infected WT mice (**Suppl. Fig. 5A**). we have previously shown that there is no difference in intestinal permeability between GzmA KO and WT mice^36^. Surprisingly, GzmB KO mice were significantly protected against *Salmonella* infection, with no systemic bacteria found in many of the mice 5 days post infection, and no weight loss (**Fig. 4C-D**). No difference in intestinal permeability was seen in GzmB KO mice (**Suppl. Fig. 5C**). GzmB KO also had significantly lower levels of serum cytokines after infection as compared to infected WT littermate controls (**Suppl. Fig. 5B**), commensurate with the low levels of systemic infection. Thus, neither inflammatory cytokines nor intestinal permeability could explain the reduced susceptibility of GzmB KO mice to infection.

**Figure 4.**
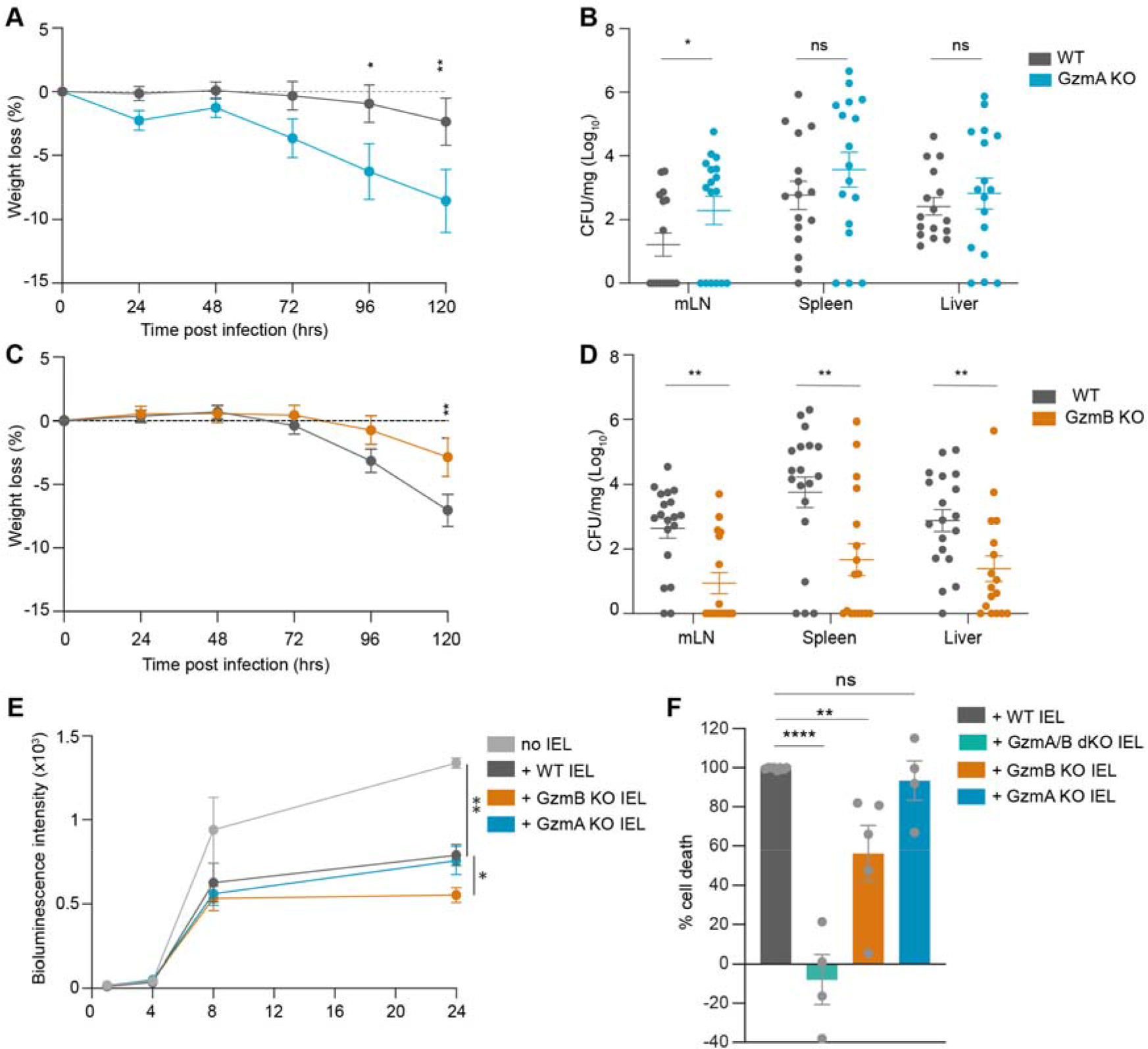
**Divergent roles of GzmA and GzmB in intestinal infection. A-B**. GzmA KO mice (n=18) and their WT littermate controls (n=19) were orally infected with SL1344-GFP and culled 5dpi. Weight loss (A) and CFU/mg in mLN, spleen and liver (B) are shown. Data were pooled from 2 independent experiments. **C-D.** GzmB KO mice (n=17) and their littermate WT controls (n=19) were orally infected with SL1344-GFP and culled 5dpi. Weight loss (C) and CFU/mg in mLN, spleen and liver (D) are displayed. Data were pooled from 2 independent experiments. **E.** MODE-K cells were infected with SL1334-lux for 1h and then incubated with GzmA or GzmB KO or their WT littermate control IEL (n=3). Bioluminescence intensity was measured when IEL were added, then after 4h, 8h and 24h. **F.** Infected MODE-K cells were stained with Crystal violet 24h after incubation with IEL from WT (n=8), GzmA/B dKO (n=4), GzmB KO (n=5), GzmA KO (n=4) mice, pooled from 5 independent experiments. The percentage cell death relative to infected MODE-K cells without IEL was normalized to the cell death induced by WT IEL. All data are presented as mean ± SEM. All data are presented as mean ± SEM. P values were calculated for bacterial counts (B, D) using Mann-Whitney U-test to compare ranks, and for all other comparisons, two-way ANOVA was used, with Sidak’s multiple comparisons tests. Standard annotations were used to denote significance: ns: not significant, * p<0.05, ** p<0.01, *** p<0.001, **** p<0.0001.

We next utilized the previously described *in vitro* infection system (**Suppl. Fig. 3C** to determine the efficacy of GzmA KO and GzmB KO IEL in controlling intracellular bacterial growth. We found that similar to WT, GzmA KO IEL were able to block the intracellular growth of SL1344-lux in intestinal epithelial cells **(Fig. 4E)**. In contrast, GzmB KO IEL were even more efficient than WT IEL, recapitulating the *in vivo* findings of GzmB KO mice infected orally with *Salmonella*. Cell viability measurements indicated that both GzmA KO IEL and GzmB KO IEL killed infected MODE-K cells much more efficiently than GzmA/B dKO IEL (**Fig. 4F**). Neither single KO recapitulated the lack of cell death, or the increased susceptibility to infection observed with the dKO, either *in vivo* or *in vitro*. Hence, we conclude that only the cell death induced by the concerted action of both Gzms is sufficient to protect against *Salmonella* infection of the intestinal epithelium.

### Absence of Granzyme A unleashes Granzyme B mediated apoptosis and luminal *Salmonella* growth

The observation that IEL lacking GzmB are better at blocking *Salmonella* growth prompted us to explore the reason for this further. Since both Gzms could induce cell death *in vitro*, we wondered if differential death modalities induced by GzmA versus GzmB might be regulating bacterial growth. A recent study showed that Salmonella utilizes nutrients from apoptotic cells to grow, thus benefiting from the induction of epithelial cell death^43^. In the presence of increased apoptosis, but not other forms of cell death, ileal *Salmonella* burdens were higher after oral infection. Of note, the increased extracellular growth of *Salmonella* in the presence of apoptotic cells was found to be dependent on a metabolic enzyme, *pflB*, a pyruvate formate-lyase, that allows Salmonella to use pyruvate from apoptotic cells as an energy source^43^. Since GzmB specifically induces apoptosis^11^, it is plausible that GzmB-induced epithelial cell apoptosis promotes growth of *Salmonella*. To test if GzmB-mediated apoptotic nutrients are also an important factor in oral *Salmonella* infection *in vivo*, we used the competitive infection system as described previously^43^. We used the Δ*pflB* strain of *Salmonella* in competition with WT *Salmonella* SL1344, to infect WT, GzmA KO and GzmB KO mice **(Fig. 5A**). The ratio of WT to Δ*pflB* bacterial counts in tissues after *in vivo* infection normalized to the input ratio (competitive index) was then analyzed in each mouse strain. Intriguingly, in the feces and ileum of GzmA KO mice, WT SL1344 had a strong competitive advantage over Δ*pflB* SL1344, which we attribute to the predominant induction of epithelial apoptosis by GzmB in these mice (**Fig. 5B-C**). In the absence of GzmB and reduced induction of apoptosis, WT SL1344 failed to exhibit a growth advantage over Δ*pflB* SL1344.

**Figure 5.**
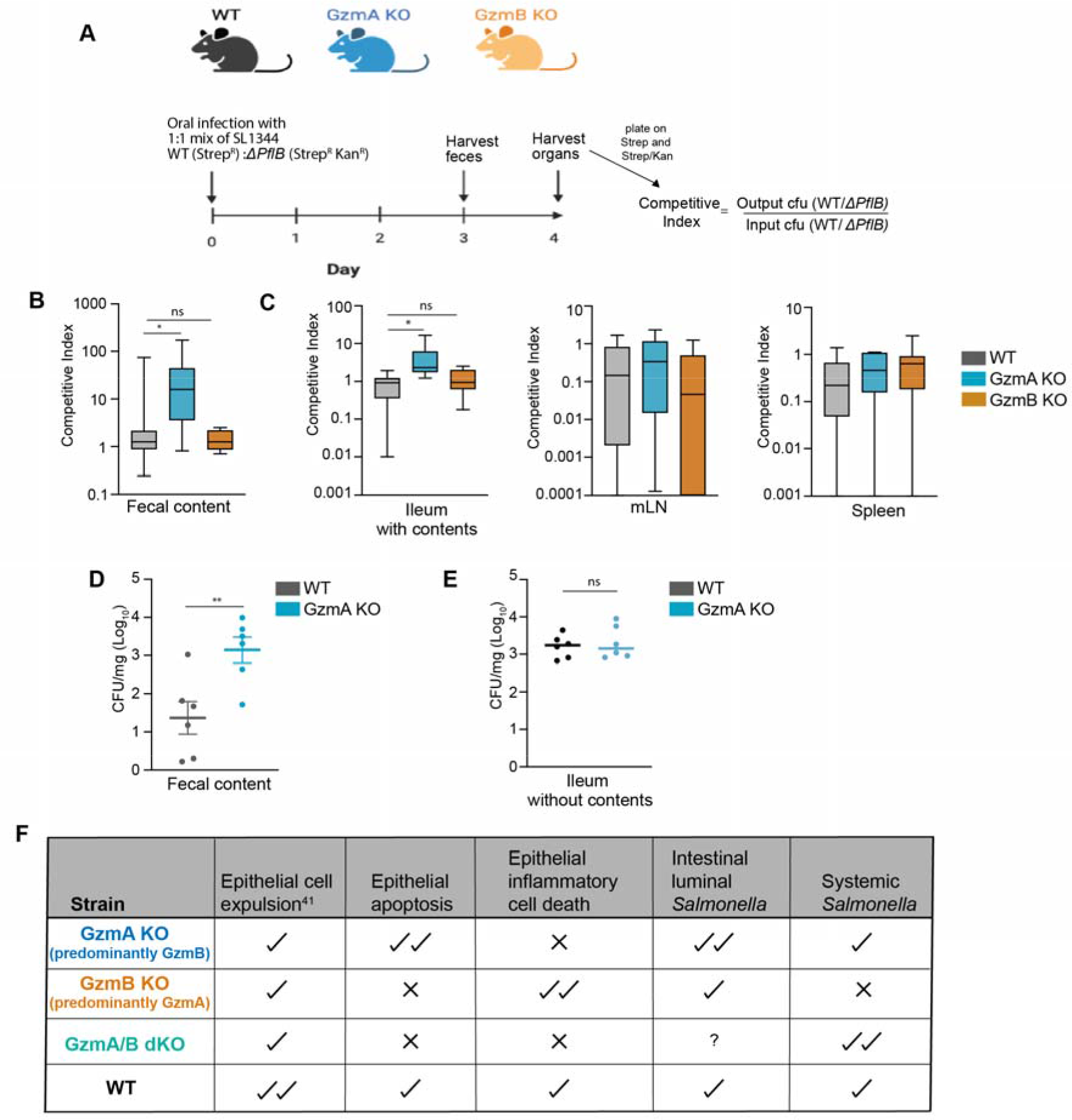
Differential cell death pathways induced by GzmA and GzmB drive divergent outcomes in infection. A. Schematic showing the set-up of competitive index experiment. WT, GzmA KO and GzmB KO generated from same line were orally infected with 1:1 mixture of wildtype: Δ*PflB* SL1344. Feces were collected at day 3 and other organs were collected at day 4 post infection. **B-C.** Box plots showing the competitive index in (B) Feces and (C) in Ileum, mLN, liver and spleen. **D-E.** WT (n=6) and GzmA KO (n=6) mice were orally infected with SL1344-GFP and culled at various days post infection (dpi) with (D) showing the fecal bacterial counts at day 3 post infection and (E) showing the bacterial counts at day 4 post infection in ileum washed with gentamycin for 30 mins to kill off any extracellular bacteria. All data are presented as mean ± SEM. For bacterial counts, ranks were compared using the Mann-Whitney U-test. Standard annotations were used to denote significance: ns: not significant, * p<0.05, ** p<0.01. **F.** Summary describing the roles of IEL intrinsic granzymes driving the phenotypes of the KO mice in intestinal *Salmonella* infection.

In GzmA KO, the competitive advantage of WT *Salmonella* was only seen at the intestinal epithelium and not in peripheral organs (**Fig. 5C**). Hence, we hypothesized that GzmB-mediated apoptosis is promoting intestinal bacterial growth in the lumen, but not translocation of bacteria across the intestinal barrier. To test this hypothesis, we enumerated bacterial loads in the feces as a measure of intestinal luminal growth. Indeed, we found that there was significantly higher bacterial burden in fecal content of GzmA KO mice as compared to WT mice (**Fig. 5D**). However, in gentamycin-treated ilea of these mice, there were similar bacterial loads, indicating similar bacterial translocation across the intestinal barrier (**Fig. 5E**). Thus, in the absence of GzmA, the balance of cell death is tipped towards more GzmB-mediated apoptosis, inducing stronger intestinal luminal growth of *Salmonella*. Overall, we conclude that the balance of cell death induced by the different Gzms provides an extra layer of complexity in the protection of the intestinal epithelium from *Salmonella* infection.

## DISCUSSION

In this study, we explored the relative contributions of GzmA and GzmB in providing protection against infection at the intestinal barrier. To our knowledge, this is the first study showing a direct protective effect of IEL-derived Gzms in intestinal infection. We find that granzyme-mediated protection was independent of perforin, and was more complex than anticipated, as it appears to depend on the concerted, but not redundant, actions of GzmA and GzmB. We show that IEL respond strongly to intestinal Salmonella infection by degranulation. In the absence of both GzmA and GzmB, IEL no longer degranulate, and are defective in killing infected epithelial cells. Moreover, Gzms may contribute to the expulsion of infected epithelial cells, as LPS-induced intestinal epithelial cell shedding requires perforin-independent activities of both GzmA and GzmB^41^. The observation of fewer cleaved-caspase-3-positive enterocytes in GzmA/B dKO mice relative to WT following TNF exposure, suggests that Gzms may also facilitate cell-intrinsic epithelial apoptosis^41^. Intriguingly, we find that the absence of Gzms also reduced the amount of pyroptosis induced by NLRC4 activation in the intestine, suggesting that Gzms aid in epithelial cell-intrinsic responses to pathogen triggers. Thus, in the absence of GzmA and GzmB, reduced epithelial cell expulsion and cell death lead to higher bacterial translocation and systemic infection with *Salmonella* (**Fig. 5F**).

Surprisingly, we found that in the absence of GzmB alone, mice were less susceptible to infection. This seemingly counter-intuitive finding could be explained by the fact that in the absence of GzmB, GzmA is the predominant granzyme, which drives lytic and possibly inflammatory cell death^44^. GzmA-mediated cell death is very effective in controlling epithelial *Salmonella* infection both *in vitro* and *in vivo* (**Fig. 5F**). In GzmA KO, induction of epithelial apoptosis by GzmB supports the growth of *Salmonella* in the ilea of infected mice. Cell death induced by GzmA (in GzmB KO) did not support the growth of *Salmonella,* consistent with its inability to induce apoptosis of target cells. Despite increased luminal growth in orally infected GzmA KO mice, only a mild increase in systemic bacterial load was observed. Here, predominant GzmB-induced apoptosis is still sufficient to clear local infection^45^, and through Casp7 cleavage, mediate extrusion of *Salmonella* infected epithelial cells^46^, which together limit bacterial translocation (**Fig. 5F**). Although we had expected that wildtype *Salmonella* would have a growth advantage over the Δ*PflB* strain in WT mice, we found that both strains grew equivalently in the intestines of WT mice, similar to previous findings^43^. It is possible that the cell death induced by the synergistic actions of GzmA and GzmB in WT mice is neither apoptotic nor inflammatory, but a combination of multiple cell death pathways that does not release nutrients that can be utilized by *Salmonella. Salmonella* infection itself induces cell-intrinsic apoptosis, pyroptosis, and panoptosis that contribute to protection of the host, but can also be co-opted by *Salmonella* for its own benefit^6^. Our data indicate that IEL intrinsic Gzms provide another layer of control for the host to optimally control *Salmonella* infection by contributing to multiple cell death pathways and expulsion. This additional layer of Gzm mediated cell death can prevent the evasion of pathogens that have evolved to modulate and benefit from intrinsic cell death pathways.

Divergent roles of GzmA and GzmB have been observed previously. In rodent lung filarial infection, GzmB KO mice were more resistant to infection, whereas GzmA KO had increased worm burdens as compared to WT mice^47^. These opposite phenotypes were ascribed to the pro-inflammatory functions of GzmA, and the anti-inflammatory functions of GzmB. In addition to infection-induced inflammation, it has been reported that the anti-tumoral advantage observed in mice that do not express GzmB is lost when, additionally, GzmA is absent in GzmA/B dKO mice^48^. These data support the idea that GzmA and GzmB are not redundant but are rather key modulators of the immune response.

The route that pathogens use to infect their hosts has a profound impact on the evolution of the host’s immune adaptation. Evolution of Gzms in particular, show evidence of species-specific adaptation to selection pressures enforced by the diversity of pathogens and the environment^7^. Constitutive expression of Gzms in intraepithelial lymphocytes provides an additional layer of protection against diverse pathogens that target barrier sites. Gzms play multi-faceted roles that involve novel activities that are still being assigned, such as the function of human GzmA in driving pyroptosis by cleaving Gasdermin B^44^. While it remains to be established exactly how GzmA functions in *Salmonella* infection, especially since the activity of this protease is enhanced by GzmB, we have now established that GzmA protects against intestinal infection. Importantly, our work establishes a new paradigm in which a perforin-independent role of GzmA and GzmB mediates protection against enteric *Salmonella* infection, and provides a rational explanation for the high levels of extracellular GzmA seen in patients with typhoid fever^14^.

## Funding and acknowledgements

The authors would like to thank the staff of the Biological Resource units, the MRC Genotyping team, the Flow Cytometry and Imaging facilities from the University of Dundee, and the animal care staff and Servicios Científico Técnicos del CIBA (IACS-University of Zaragoza) and Servicio Apoyo Investigación (University of Zaragoza) for invaluable assistance. This work was supported by the Wellcome Trust and Royal Society (Sir Henry Dale Fellowship to MS, 206246/Z/17/Z) and, in part by a Wellcome Trust Senior Investigator award [106914/Z/15/Z] to JCDH, National Institutes of Health grant R01AI170617 and R01DK135272 to KLE, by ARAID Foundation and grant SAF2017-83120-C2-1-R and PID2020-113963RBI00 from the Ministry of Science and Innovation/ Agencia Estatal de Investigacion and FEDER (Group B29_17R, Aragon Government) to JP. LS was supported by a PhD fellowship (FPI) from the Ministry of Science, Innovation and Universities. NS is supported by a Medical Research Scotland PhD studentship. MA was supported by a post- doctoral fellowship “Juan de la Cierva-formación” and “Juan de la Cierva-incorporación” from the Ministry of Science, Innovation and Universities. NBG is supported by New Jersey Commission on Cancer Research Predoctoral Fellowship COCR23PRF032. We would also like to acknowledge Prof. Peter Vandenabeele and Dr. Wei Xie from VIB, University of Gent, for helpful discussions and reagents.

For open access, the authors have applied a CC BY public copyright licence to any Author Accepted Manuscript version arising from this submission.

## Author contributions

MS conceived of the study. MV, ASC, JP and MS conceptualized and designed experiments. ASC, MV, MA, LS, SJT, DD, CN, MGT, NBG, KLE and MS performed experiments. EO, MB, NW, JCDH, CJA made critical bacterial strains. ASC, MV and MS drafted the manuscript with substantial input from KLE, CJA, JP and JCDH. All authors contributed to revising and critically reviewing the manuscript and approved the final manuscript.

## METHODS

### Mice

*GzmA^-/-^GzmB^-/-^, GzmA^-/-^, GzmB^-/-^* were bred in the University of Dundee and *Prf1^-/-^* (Perforin) mice were bred at the University of Zaragoza. *GzmA^-/-^GzmB^-/-^* and *Prf1^-/-^* mice were randomly co-housed with control age- and sex-matched C57BL/6J mice at weaning (between 3-4 weeks of age), whereas for experiments with *GzmA^-/-^* or *GzmB^-/-^* mice, wild-type littermate controls were used wherever possible. C57BL/6J and *Rag2^-/-^*(RAGN12) mice were purchased from Charles Rivers and Taconic, respectively. Animals were housed in a standard barrier facility at 21°C, 55-65% relative humidity with a 12h light/dark cycle. Mice were given *ad libitum* access to water and a diet of RM1 (autoclaved RM3 when in breeding). Environmental enrichment of plastic tubes, play houses and additional sizzle nest bedding were given to all cages. All mice were bred and maintained with approval by the University of Dundee Welfare and Ethical Use of Animals committee, under a UK Home Office project license (PD4D8EFEF) in compliance with U.K. Home Office Animals (Scientific Procedures) Act 1986 guidelines, in an Association of the Assessment and Accreditation of Laboratory Animal Care-accredited facility according to protocols approved by the Center for Comparative Medicine and Surgery at Icahn School of Medicine at Mount Sinai, or approval by the University of Zaragoza ethical review committee (PI53/20) designated as Animal Welfare Body (Article 34, Royal Decree 53/2013). In addition, a local study plan was approved as legally compliant detailing the animals involved, the experimental procedure(s), local scoring system, potential adverse effects and humane endpoints, and the named responsible PIL(s).

### Experimental Design

Sample size was determined based on the CFU in MLNs of infected WT mice, and power calculations based on a one-sided type I error of 0.05, aiming to achieve statistical power of 0.80 to find effect size of at least 1 standard deviation. For each experiment, WT animals were compared to age and sex matched gene-deficient mice. Mice were randomly assigned to cages co-housed at weaning to prevent “cage effects” and acclimatised for at least 10 days prior to dosing. Since all mice were infected in this study, no further randomisation was done. Dosing was performed in sequence with the tissue harvest following the same sequence. Dosing, experimental monitoring and analysis were performed in a blinded fashion. No mice were excluded other than those that had to be culled within 24 hours of oral dosing because they reached the severity limit as predefined in the study plan, suggesting that systemic infection had occurred.

### Bacterial strains

Derivatives of *Salmonella enterica* serovar Typhimurium strain SL1344 were used: SL1344-GFP, containing a plasmid expressing GFP-Ova (pMW57) under the control of the pagC promoter (obtained from D. McEwan, Dundee with permission from D. Bumann, Basel) ^49^; SL1344-Lux was generated by introducing pEM7-lux plasmid (that expresses l*uxCDABE* constitutively under the control of the synthetic promoter PEM7 ^50^); ΔSPI1 strain (JVS-00405) was obtained from J. Vogel’s lab (Würzburg), and is a λ-Red knockout of the entire SPI1 island ^51^, SL1344 WT and ΔpflB were obtained from C.J Anderson^44^.

### Infections, CFU determination and competitive index

9-12 weeks old female (18g-23g) and male (24g-30g) mice were infected with ampicillin-resistant SL1344-GFP. Bacteria were grown overnight in LB broth supplemented with ampicillin (100 μg/ml) at 37°C, on a shaker, then subcultured in LB+ ampicillin medium for at least 3h before in vivo infection. Bacteria were centrifuged 10 min at 3750 rpm, washed and resuspended in sterile PBS. The OD600 was measured to estimate bacterial density. Serial plating on LB agar supplemented with ampicillin (100 μg/ml) were performed to quantify the infection dose. For oral infection, food was removed at least 3h before gavage and mice were infected with 1.5 x 10^8^ bacteria. For intravenous infection, mice were infected with 500 bacteria. On the day of infection, food was removed at least 3h before gavage. Mice were orally infected with 100μl of PBS containing a total of 10^7^ bacteria. To determine if epithelial cell invasion is necessary for granzyme action, the ΔSPI1 strain was used at a concentration of 9 x 10^9^ bacteria per mouse. In addition to weight loss monitoring, mice were monitored a minimum of twice a day for apparition of clinical signs and a scoring system with defined humane endpoints was used to assess the severity of infection. For competitive index assays, WT (Strep^R^) and Δ*PflB* (Strep^R^ +Kan^R^) SL1344 strain were mixed at 1:1 ratio in 100 μl of PBS containing 1 x 10^9^ bacteria/strain. The input ratio of each strain was calculated by plating the infective dose on agar plates containing streptomycin (total inoculum) and agar plates containing streptomycin plus kanamycin (Δ*PflB* strain). Wild-type *Salmonella* input cfu was calculated by = cfu of total inoculum−cfu of Δ*PflB*, and the input ratio was calculated as cfu(WT)/cfu(Δ*PflB*).

For output cfu determination, mice were euthanized by a rising concentration of CO_2_ and death was confirmed by cervical dislocation. Fecal content, Ileum, caecum, colon, mLNs and spleen were collected, weighed, and transferred into 2ml Precellys (Bertin) tubes containing ceramic beads and PBS + 0.05% Triton X-100 (Sigma). Tissues were homogenized using the Precellys 24 homogenizer (Bertin) for 2x10s at 5000Hz. Livers were weighed and crushed through a 70μm strainer in PBS+ 0.05% Triton X-100. Supernatants were serial diluted in LB + ampicillin (100μg/ml) medium and plated on LB + ampicillin plates. Colonies were counted after overnight incubation at 37°C. For quantification of bacteria in competitive index assay, supernatants from organs were plated on LB plates containing Streptomycin and Streptomycin + Kanamycin. Output cfu ratio was calculated as cfu(WT)/cfu(Δ*PflB*). Competitive index was calculated as Output/input ratios.

### FlaTox treatment and pyroptosis imaging

WT and GzmA/B dKO mice^41^ bred at Icahn School of Medicine at Mount Sinai were administered FlaTox (0.16µg/g PA (Protective Antigen) and 0.08µg/g LFn-Fla (the fusion of flagellin protein FlaA and N-terminal domain of anthrax lethal factor (LFn)), generously provided by Isabella Rauch, OHSU) intravenously for 60 min. *In vivo* propidium iodide staining was performed by injecting mice with 100μg intravenously 10min prior to euthanasia. Jejunal tissues were fixed in PLP buffer (0.05M phosphate buffer containing 0.1M L-lysine [pH 7.4], 2 mg/ml NaIO_4_ and 1% PFA) overnight as previously described^40^. Tissues were embedded in OCT, sectioned at 5µm, stained with Hoechst 33342 and AlexaFluor 647-phalloidin (Invitrogen), and mounted with ProLong Glass (Invitrogen). Images were acquired on an inverted Ti2 microscope (Nikon) equipped with a Dragonfly spinning disk (Andor), Sona sCMOS camera (Andor), PLAN APO 20x/0.8NA dry objective, and Fusion acquisition software (Andor). The number of PI^+^ cells per 0.1 mm^2^ villus was quantified by an observer blinded to the condition.

### *In vivo* intestinal permeability assay

Naïve and infected mice were treated with FITC-dextran dissolved in PBS (100mg/ml). Prior to treatment, mice were starved for 4h. Each mouse received 44mg of FITC-dextran solution per 100g body weight by oral gavage. After 4h, blood was collected, and the serum was separated. The concentration of FITC-dextran in serum was quantified by spectrophotometry with an excitation of 485 nm using a CLARIOstar Plus plate reader.

### IEL isolation and culture

IEL were isolated as described by James *et al*^52^. Briefly, small intestines were cut from proximal duodenum to terminal ileum and flushed with 20 ml of cold PBS. Small intestines were longitudinally opened, then transversely cut into ∼5 mm pieces and put into 25 ml of warm RPMI medium (RPMI; 10% FBS; 1% glutamine; 1% penicillin-streptomycin), containing 1mM DTT. Small intestine pieces were agitated on a rotator or bacterial shaker for 40min at room temperature, centrifuged, vortexed and passed through a 100μm sieve. Cells were centrifuged in a 36%/67% Percoll/PBS density gradient at 700g for 30 minutes. Total ΙEL were collected from the interface between 36% and 67% Percoll. CD8α+ IEL were purified using the EasySep mouse CD8α positive selection kit II (STEMCELL Technologies), with small changes from the manufacturer’s instructions: Total IEL were resuspended in 250μl of isolation medium. After incubation with Fc blocker and antibody cocktail mix (at a concentration of 10μl/ml and 50μl/ml, respectively), 80μl/ml of dextran beads were added to the cells. Labelled cells were then incubated in the magnet for 5min before pouring off the supernatant. The remaining cells were collected and used as CD8α+ IEL. Purity ranged between 75%-85%.

For *in vitro* degranulation assay, isolated IEL were incubated in 100ng/ml of IL-15/Ra complex (Thermo Fisher Scientific) overnight at a concentration of 1 million cells per well/ml in a 24 well plate. Next day, cells were washed once, and resuspended in 100ng/ml IL-15 with 50ng/ml of PMA +1μg/ml of ionomycin (Sigma) at the density of 0.5 million cells/well. After 30 minutes, anti- CD107a (clone 1D4B, Biolegend) and CD107b (clone M3/84Biolegend), Brefeldin (Biolegend) and Golgi stop (BD Biosciences) were added. Post 4 hrs of culture, cells were washed and fixed analyzed for CD107 intensity by flow cytometry.

### LPL isolation

Small intestines were cut from proximal duodenum to terminal ileum and flushed with 20 ml of cold PBS. Small intestines were longitudinally opened and stored on ice in 10 ml of PBS. Samples were vortexed 3 times for 10 seconds in PBS. Small intestines were then incubated for 30 min, with constant shaking, in “strip buffer” (PBS; 5% FBS; 1mM EDTA, 1mM DTT), prewarmed at 37°C. After the incubation, tissue was washed in PBS, then incubated for 45 min with constant shaking with “digest buffer” (RPMI; 10% FBS; 1mg/ml collagenase/dispase (Roche); 20μg/ml DNAse 1 (Sigma)). Supernatants containing LPL were then collected by pouring over a 70μm filter into a new tube containing RPMI media supplemented with 10% FBS.

### Adoptive transfer

6 to 8 weeks old male and female *Rag2^-/-^* mice were intravenously injected with CD8α-sorted IEL, previously cultured for 24h with IL-15/IL-15R complex recombinant protein (100ng/ml, Thermo Fisher Scientific) and retinoic acid (100ng/ml). The efficiency of IEL transfer was assessed 4 weeks post transfer by flow cytometry. For IEL competitive adoptive transfer, mice were injected with a mix of WT and GzmA/B dKO IEL (1x10^6^ cells in 100ul of PBS; 1:1 ratio). For *Salmonella* challenge experiments, *Rag2^-/-^*mice were transferred with 10^6^ WT or GzmA/B dKO IEL and 4 weeks later mice were orally challenged with SL1344-GFP as described above.

### Flow cytometry

The following murine monoclonal antibodies were used to detect cell surface markers: TCRβ [clone H57-597 (BioLegend)], TCRγδ [clone GL3 (BioLegend or eBioscience)], CD4 [clone RM4-5 (BioLegend)], CD8α [clone 53-6.7 (BioLegend)], CD8β [clone H35-17.2 (eBioscience)], CD103 [clone 2E7 (BioLegend)], CD160 [clone 7H1 (BioLegend)],CD223 (LAG3) [clone eBioC9B7W (eBioscience)], CD85k (LILRB4) [clone H1.1 (BioLegend)], TIGIT [clone GIGD7 (BioLegend)], CD69 [clone H1.2F3 (eBioscience)], CD96 [clone 3.3 (BioLegend)], CD45 [clone 30-F11 (BioLegend)], Nkp46 [clone 29A1.4 (BioLegend)]. For intracellular staining, cells were fixed with 2% PFA at 37°C for 10 min before permeabilization with permabilization buffer (eBioscience). Cells were incubated with the following murine monoclonal antibodies: GzmB [clone GB12 (eBioscience)], GzmA [clone GzA-3G8.5 (eBioscience)], FoxP3 [clone FJK-16s (eBioscience)].

### mLN and spleen cell suspension

Mice were culled by CO_2_ and tissues were collected. mLN were crushed through a 70μm strainer in RPMI medium (RPMI; 10% FBS; 1% glutamine; 1% penicillin-streptomycin). Spleens were crushed through a 70μm strainer and red blood cells were lysed, and remaining cells taken for flow cytometric analyses.

### K562 killing assay

For testing the cytotoxicity of WT and GzmA/B dKO IEL, a bioluminescence based cytotoxicity assay was used as previously described^53^. Luciferase-expressing K562 cells (kind gift of Dr. S. Minguet, Freiburg) were plated at 5x10^3^ cells/well in a 96-well plates. 75μg/mL D-firefly luciferin potassium salt (Biosynth) was added to the cells and bioluminescence was measured with a PHERAstar plate reader to establish the bioluminescence baseline. For a maximal cell death (positive control), Triton X-100 was added to K562 cells at a final concentration of 1%. For spontaneous death (negative control), culture medium was added to K562 cells. For the test, WT or GzmA/B dKO IEL that had been cultured overnight in 20ng/mL IL-15 (Peprotech) were added to K562 cells at a 40:1 effector-to-target (E:T) ratio with anti-CD3 (1μg/ml) and incubated for 24h. Bioluminescence was measured as relative light units (RLU). Percentage specific lysis was calculated as follows: % specific lysis = 100 x (average spontaneous death RLU – test RLU) / (average spontaneous death RLU – average maximal death RLU).

### MODE-K infection and CFU assays

Ampicillin-resistant SL1344-Lux were grown as described in the above sections and were resuspended in infection medium (DMEM + 1% BSA, without antibiotics). MODE-K (kind gift of. D. Kaiserlian, Lyon) were seeded overnight (in DMEM supplemented with 10% FBS and 1% L-glutamine, no antibiotics) and infected at a multiplicity of infection (MOI) of 10. For intracellular growth assessment, bacteria were pre-mixed with CD8α^+^ purified IEL (1:10 bacteria-effector ratio), or left untreated, for 15 min and the mix was then added to the MODE-K, for 1h. Cells were washed and treated with 50μg/ml of gentamicin in infection medium to kill off extracellular bacteria for 30 min (90 min time point) and then kept in 12.5μg/ml of gentamicin. At indicated time points, bioluminescence was measured in a PHERAstar plate reader. To assess bacteria extracellular growth in presence of IEL, SL1344-lux were mixed with IEL (1:10 bacteria-effector ratio) and bioluminescence was measured at indicated time point. For assessment of intracellular growth of *Salmonella*, MODE-K cells were first infected with SL1344-lux (MOI of 10) for 1h, washed and treated with 50μg/ml of gentamicin. Then CD8α sorted IEL (1:10 bacteria-effector ratio) along with 100ng/ml of IL-15/IL-15R complex recombinant protein were added. At indicated time points, bioluminescence was measured using a PHERAstar plate reader.

After 24h, survival of the MODE-K cells was assessed, by first washing the cells in PBS, to remove IEL and bacteria, then staining living cells for 20 minutes with 100μl/well of Crystal Violet (0.75g in 12.5ml water and 37.5ml methanol). After washing with water, the Crystal Violet was solubilised by adding 200μl/well of methanol. The solubilised Crystal Violet was quantified by reading absorbance at 570nm. The absorbance was normalised to the absorbance in infected MODE-K cells without IEL, to calculate the percentage cell death, using the formula : cell death (%)=100* [Abs(MODE- K_inf_) – Abs(MODE-K_inf+IEL_)/Abs(MODE-K_inf_)].

### Immunofluorescence and imaging

Small intestinal tissue from GzmA sKO or GzmB sKO mouse or their WT littermates were flushed once with cold HBSS and once with a room-temperature fixative (2% paraformaldehyde in PBS, pH=7.4), and small (0.8-1 cm) fragments from jejunum or ileum were excised and incubated at room temperature in 10 ml of fresh fixative for 2-3 hours with slow agitation. The tissues were then washed three times with 50 mM ammonium chloride in PBS and incubated overnight in PBS containing 30% sucrose, before embedding in OCT (Agar Scientific) on dry ice and stored below - 18^0^C. 15 µm sections of the frozen tissues were rehydrated in PBS, permeabilised for 10-15 min with 1% NP40 in PBS, blocked for 1 hour with PBS containing 2% BSA and 0.1% Triton-X100 and stained with a mixture of Alexa Fluor 647-labelled mouse anti-Granzyme A ab (Santa Cruz, clone 3G8.5, sc33692-AF647, at 1:500 dilution) and goat anti-Granzyme B ab (R&D systems, AF1865, at 1:50 dilution) in above blocking solution overnight at +4^0^C. After 5 washes with PBS, the slides were stained for 1 hour with Alexa Fluor 647-conjugated donkey anti-mouse, Alexa Fluor 488- or Rhodamine Red-X-conjugated donkey anti-goat antibodies (Jackson ImmunoResearch, at 1:500 dilution) and Alexa Fluor 568 phalloidin (Invitrogen, at 1:75 dilution) or acti-stain 488 phalloidin (Cytoskeleton, Inc, at 1:75 dilution) in the blocking solution. After 4 additional PBS washes, the tissues were stained with 1 µg/ml DAPI in PBS for 10 min, washed in PBS and mounted using Vectashield Vibrance antifade mounting media (Vector Laboratories). Tissues were imaged using confocal Zeiss LSM 710 or LSM 880 microscope operated by Zen software using 63x/1.4NA oil immersion objective. Where indicated, serial optical sections were collected throughout the entire thickness of the tissue, and maximal intensity projections from the sections spanning selected IELs were generated in ImageJ.

### Plasma preparation and cytokine level measurement

Mice were euthanized by CO_2._ Blood was collected by intracardiac puncture and transferred into tubes containing 5mM EDTA. Blood was centrifuged for 15 min at 800*g*, and plasma was collected. Plasma cytokine levels were assessed using mouse ProcartaPlex cytokine and chemokine panels (ThermoFisher Scientific), according to the manufacturer’s instructions. Data were acquired using the Luminex 200 analyser.

### IEL:enteroid co-cultures

To generate the enteroids, small intestine from a C57BL/6J mouse were longitudinally cut to scrape off the villi using a coverslip and washed with cold PBS. The small intestine was cut into small 3-5mm pieces and incubated in PBS containing 1mM EDTA for 20 minutes at 4°C on a tube roller. After being filtered through a 100µm sieve, the pieces were further incubated in PBS supplemented with 5mM EDTA for 30 minutes at 4°C. The tissue fragments were transferred into PBS and vigorously shaken for one minute. Isolated crypts were centrifuged for 3 minutes at 200*g*, mixed with Matrigel (Corning) and plated in a 24-well plate in ENR medium (Advanced DMEM/F12 supplemented with 50ng/mL EGF (PeproTech), 100ng/mL Noggin (PeproTech) and 1µg/mL R-spondin-1 (PeproTech))

For IEL-enteroid co-culture, two-day-old enteroids were released from the Matrigel and incubated with WT or GzmA/B dKO purified CD8α^+^ IELs that were labelled with cell tracking dye CFSE, for 30 minutes at 37°C (at a ratio of 500 IELs to 1 enteroid) and then plated in an 8-well μ-Slide ibiTreat (Ibidi) in ENR medium supplemented with 10ng/mL soluble IL-15 and 100 U/mL IL-2. 48h later, live imaging of the co-culture was performed using Zeiss 710 confocal microscope system at 37°C and 10% CO_2_, 20X dry objective. 20 z-stacks with 3µm interval of the region of interest (ROI) were acquired for at least 90 min. The mean track speed and track displacement length were analyzed using Imaris by creating a Spot function that detected cells with diameter from 7.5µm - 8µm.

### Statistical analysis

Data analysis was done using GraphPad Prism v9. For bacterial counts, ranks were compared using the Mann-Whitney U-test. For all other comparisons, two-way ANOVA was used, with multiple comparisons using Sidak’s multiple comparisons tests, unless otherwise stated in the figure legend. Standard annotations were used to denote significance: * p<0.05, ** p<0.01, *** p<0.001, **** p<0.0001.

### Schematic diagrams

Schematic diagrams in this manuscript were created using Biorender.

### Data availability

No large datasets were generated during this study. All data are shown in the figures, and raw data supporting this study can be obtained from the corresponding author upon request.

