## Supplemental figures 1-5 for "Distinct cell death pathways induced by granzymes collectively protect against intestinal *Salmonella* infection"

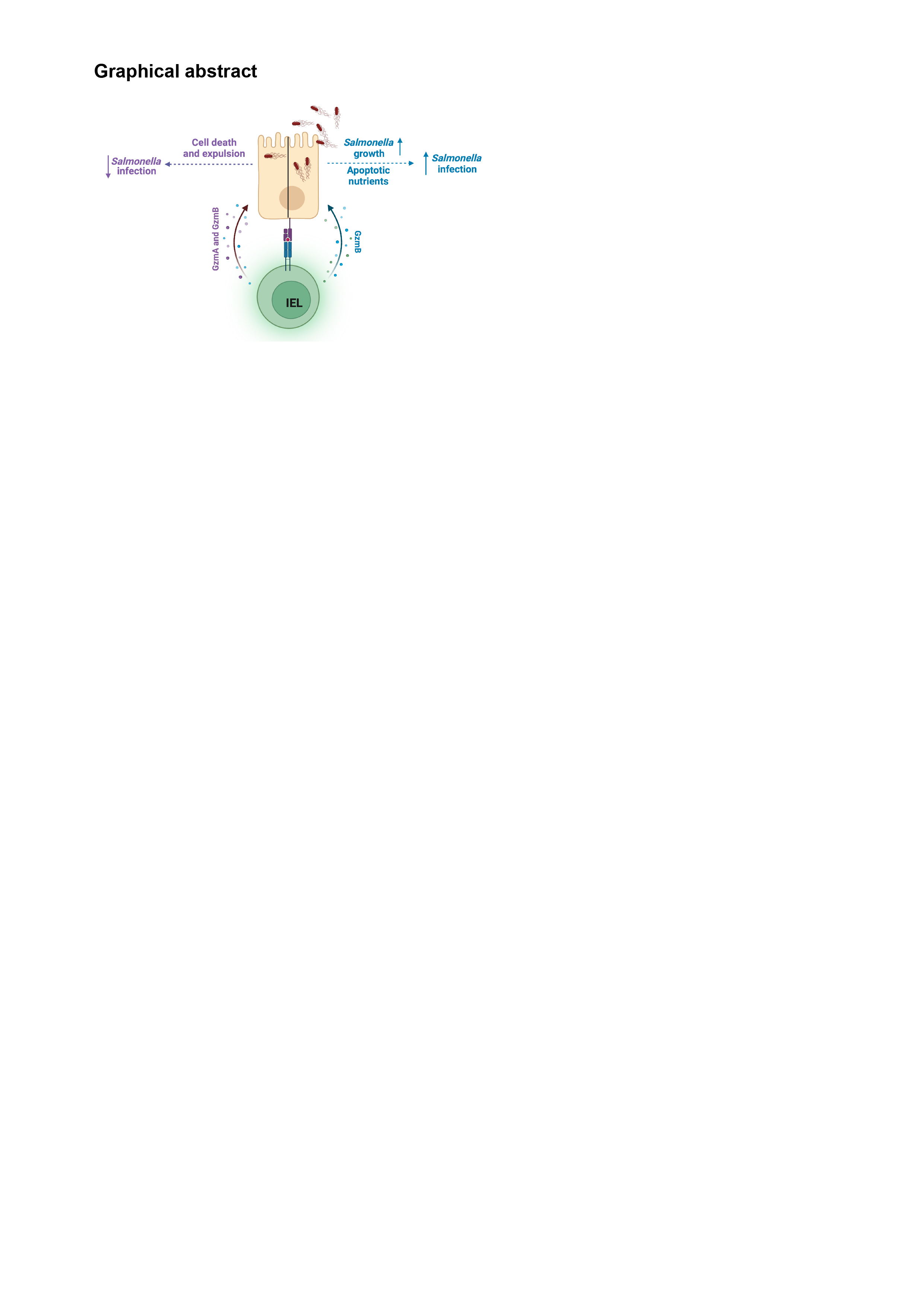

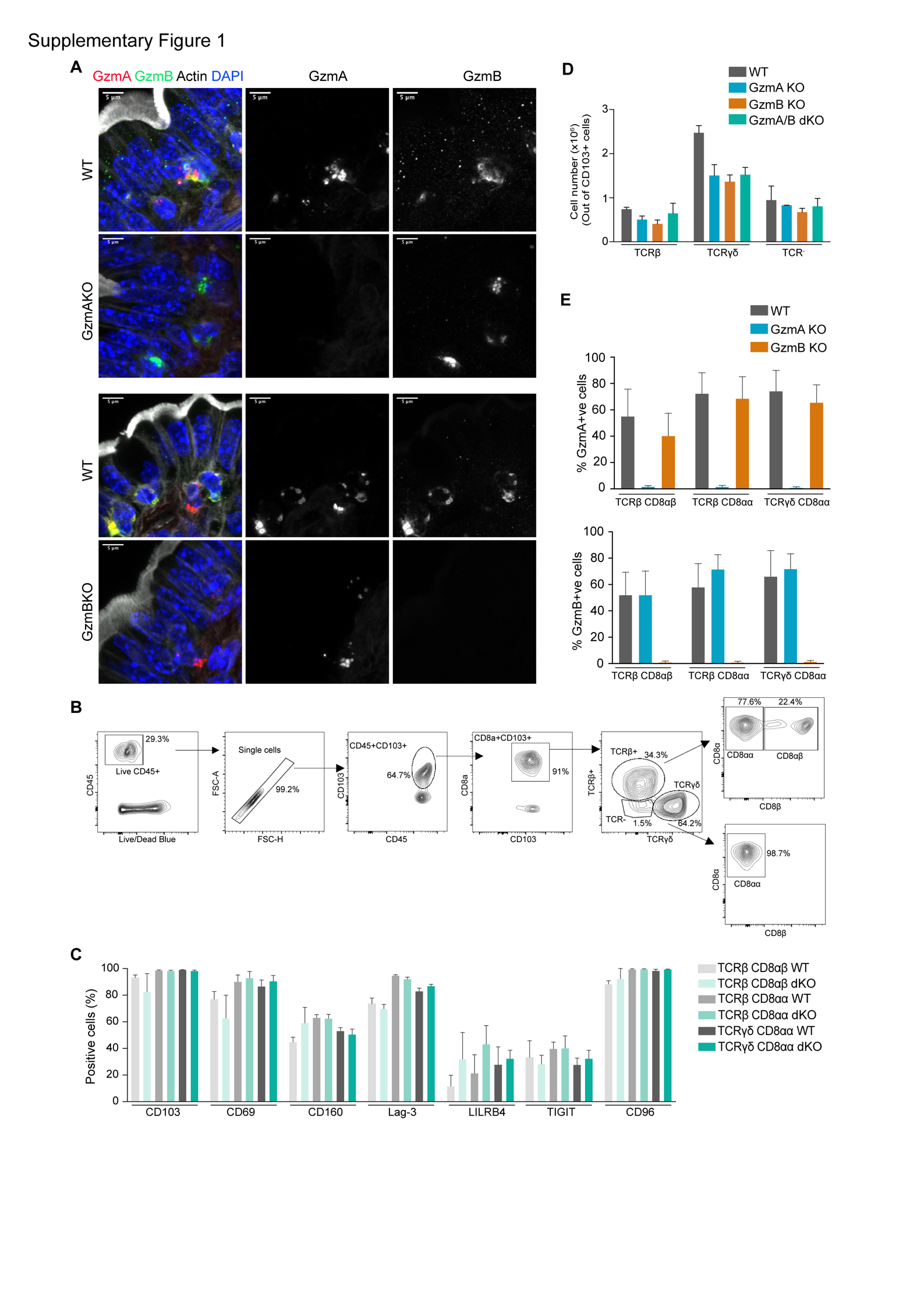
**Supplementary Figure 1**. **A.** Representative immunofluorescent micrographs to confirm the specificity of the GzmA and GzmB antibodies. Images show cells expressing GzmA (red) and GzmB (green) in the epithelial layer of a jejunal villus in WT, GzmA KO and GzmB KO. Sections were counterstained with phalloidin to show actin (white) and DAPI to show nuclei (blue on left panel). Scale bar=50µm. **B.** Gating strategy of IEL in the small intestine epithelium. Total IEL were gated as live, single, CD45^+^CD103^+^ cells, which can be further divided into TCRβ^+^, TCRγδ^+^ and TCR^–^ subsets. These subsets are mainly CD8α^+^ T cells. **C.** Comparison of the percentage of positive WT and GzmA/B dKO IEL, from co-housed mice, for CD103, CD69, CD160, Lag-3, LILRB4, TIGIT and CD96 surface markers (n=3 each). **D.** Absolute numbers of CD103^+^ IEL in WT littermate controls, GzmA sKO, GzmB sKO and GzmA/B dKO mice. **E.** Comparison of the percentage of GzmA^+^ (upper panel) or GzmB^+^ (lower panel) IEL in GzmA KO, GzmB KO, and WT mice (n>5/strain). All data are presented as mean ± SEM. P values were calculated for (C-E) by ordinary one-way ANOVA with Sidak’s multiple comparisons.  Where no p-values are shown, no significance was found.

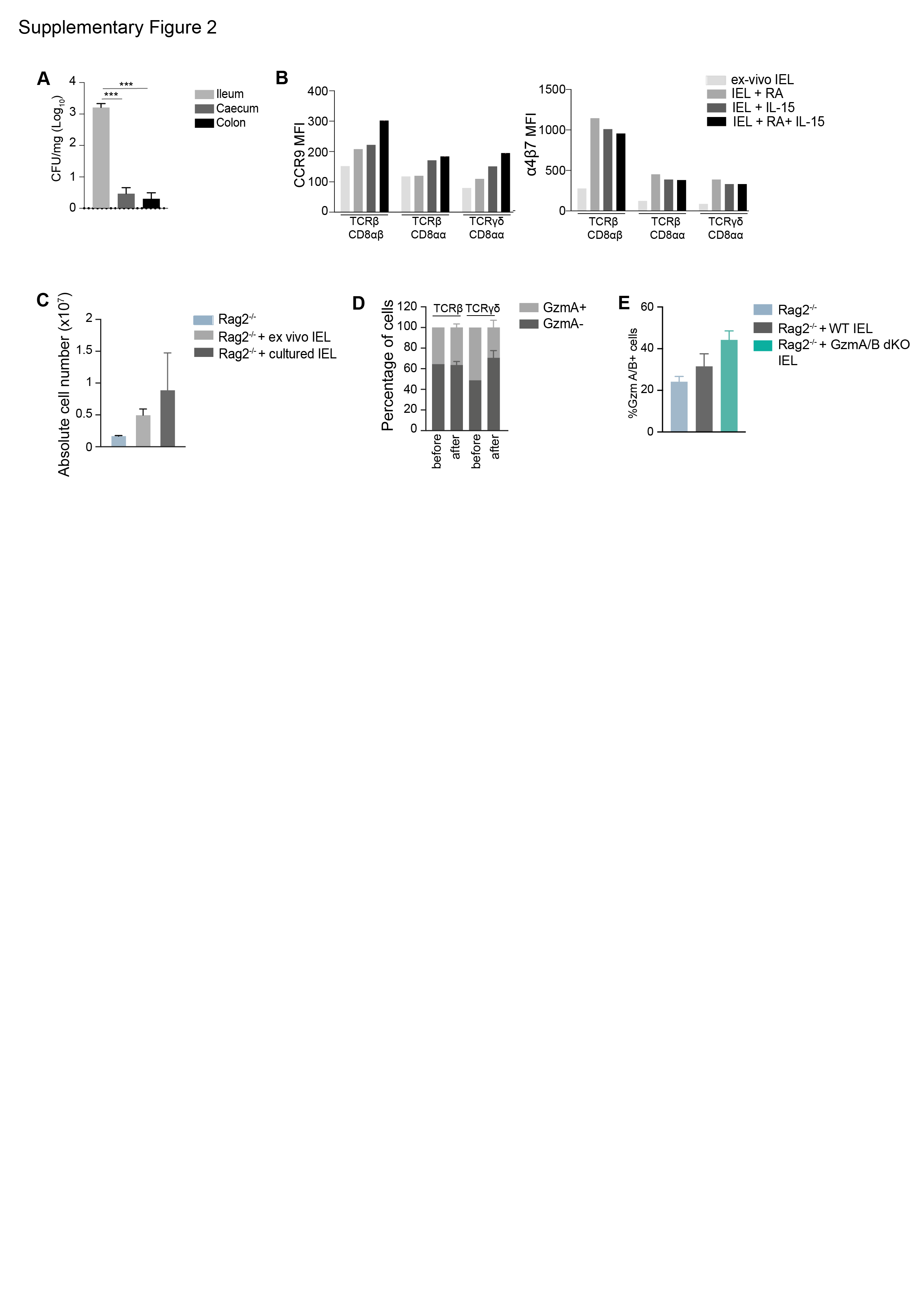

**Supplementary Figure 2**. **A.** Bar graph showing total bacterial (SL1344) counts in ileum, caecum, and colon of WT mice 4 days post infection (n=6). **B.** Comparison of the expression of CCR9 (upper panel) and α4β7 (lower panel) on ex vivo IEL and IEL cultured with retinoic acid and 100 ng/ml IL-15/Rα. **C.** Bar graph showing the number of CD45^+^ cells collected in the SI epithelium after ex-vivo (n=3) or cultured (n=2) IEL transfer into Rag2^-/-^ mice. **D.** Competitive transfer of cultured WT and GzmA/B dKO IEL into Rag2^-/-^ mice. WT and GzmA/B dKO IEL were mixed at a ratio of 1:1 before the transfer. Bar graphs show the ratio of GzmA positive cells in the TCRβ and TCRγδ populations, in the 1:1 mix before the transfer (n=2) and 4 weeks after the transfer, isolated from the adoptively transferred Rag2^-/-^ mice (n=4). All data are presented as mean ± SEM. P values were calculated for (A) using ordinary one-way ANOVA with Sidak’s multiple comparisons.  Standard annotations were used to denote significance: *** p<0.001. **E.** Percentages of GzmA+/GzmB+ cells out of CD45+ cells isolated from the small intestinal epithelia of Rag2^-/-^ mice, and from Rag2^-/-^ mice that received IEL from either WT or GzmA/B dKO mice 4 weeks earlier, and that were then infected with SL1344. These data are from the mice shown in Fig. 2G-I.

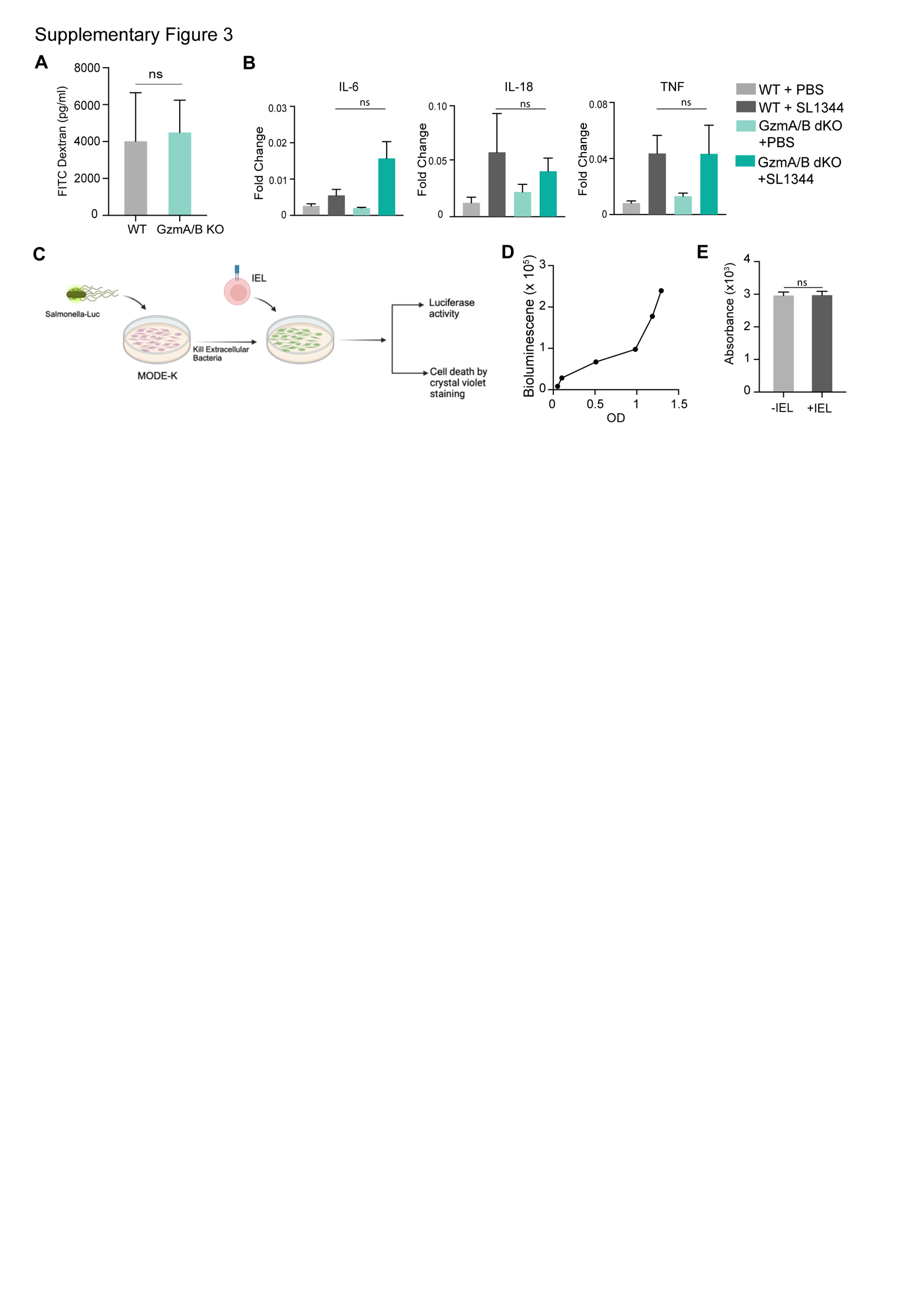
**Supplementary Figure 3**. **A.** Bar graphs comparing intestinal permeability in cohoused naïve WT and GzmA/B dKO mice (n=5/group). The concentration of FITC dextran was measured in the serum of the mice 4h after the gavage. **B.** Transcript fold change in the IL-6, IL-18 and TNF in small intestinal tissue of cohoused WT and GzmA/B dKO mice 3 days post *Salmonella* infection (n=3/group). **C.** Schematic showing the set-up of invitro MODE-K killing assay by IEL. MODE-K was infected with SL1344-lux for 0.5 h and then extracellular bacteria was killed using gentamycin. IEL were added to infected MODE-K and bioluminescence was acquired at different time points. **D.** Graph showing the correlation between bacterial OD and bacterial bioluminescence. **E.** Bar graph showing the survival, measured using crystal violet method, of uninfected MODE-K in the presence or absence of IEL (n=3/group). All data are presented as mean ± SEM. P values were calculated for (A and D) using unpaired t-test and for (B) ordinary one-way ANOVA with Sidak’s multiple comparisons. Where no p-values are shown, no significance was found. Standard annotations were used to denote significance: ns, not significant.

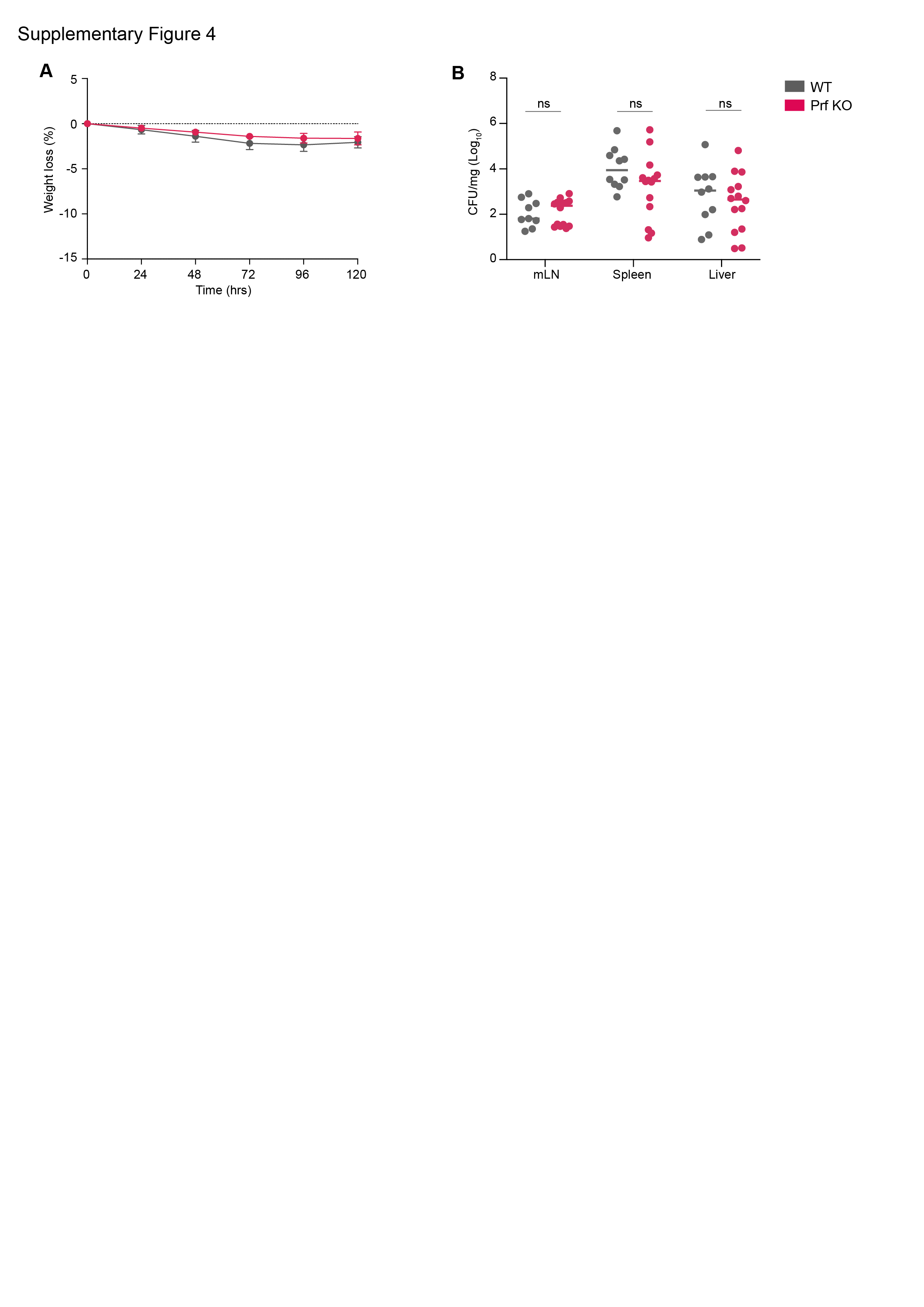

**Supplementary Figure 4. A-B.** Cohoused WT (n=10) and Pfn KO (n=14) mice were orally infected with SL1344-GFP and culled 5dpi. Weight loss (A) and CFU/mg in MLN, spleen and liver at the time of sacrifice (B) are shown. Data were pooled from 2 independent experiments. All data are presented as mean ± SEM.  P values were calculated for bacterial counts, ranks were compared using the Mann-Whitney U-test and all other comparisons, two-way ANOVA was used, with multiple comparisons using Sidak’s multiple comparisons tests. Standard annotations were used to denote significance: ns: not significant.

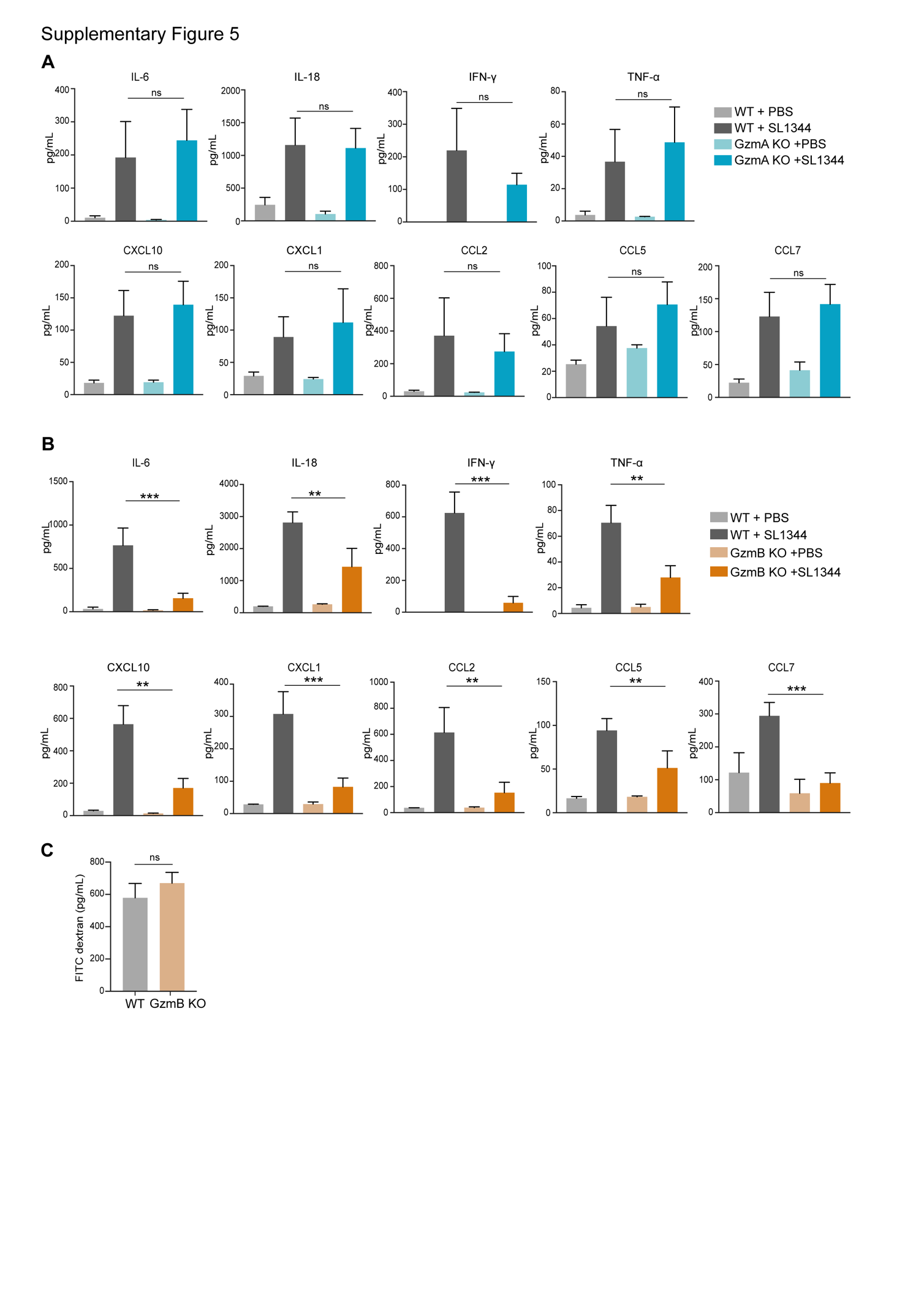
**Supplementary Figure 5.** Chemokine and cytokine levels in plasma of naïve and orally infected (**A**) GzmA sKO (n=12) and (**B**) GzmB sKO (n=12), compared to their WT littermate controls (n>9) mice from figure 4, 5 days post oral *Salmonella* infection. **C.** Dot plots comparing intestinal permeability in naive GzmB KO and WT littermate control mice (n=5/group). The concentration of the FITC dextran was measured in the serum of the mice 4h after the gavage. Data were pooled from 2 independent experiments. All data are represented as mean ± SEM. P-values were calculated for (A-B) by ordinary one-way ANOVA with Sidak’s multiple comparisons, (C) by unpaired t-test. Standard annotations were used to denote significance: ns, not significant, ** p<0.01, *** p<0.001.

**Supplemental videos 1 and 2.** Representative time-lapse confocal microscopy videos of either (**1**) WT or (**2**) GzmA/B dKO IEL (in green) co-cultured with WT enteroids. Frames were acquired every 2.5 minutes for 90 min.
